# *Pseudoalteromonas* is a novel symbiont of marine invertebrates that exhibits broad patterns of phylosymbiosis

**DOI:** 10.1101/2025.08.22.671635

**Authors:** Alejandro De Santiago, Shelby Barnes, Tiago José Pereira, Mirayana Marcellino-Barros, Lekeah Durden, Min Khant Han, J. Cameron Thrash, Holly M. Bik

## Abstract

Despite growing insights into the composition of marine invertebrate microbiomes, our understanding of their ecological and evolutionary patterns remains poor, owing to limited sampling depth and low-resolution datasets. Previous studies have provided mixed results when evaluating patterns of phylosymbiosis between marine invertebrates and marine bacteria. Here, we investigated potential animal-microbe symbioses in *Pseudoalteromonas*, an overlooked bacterial genus consistently identified as a core microbiome taxon in diverse invertebrates. Using a pangenomic analysis of 236 free-living and invertebrate-associated bacterial strains (including two new nematode-associated isolates generated in this study), we confirm that *Pseudoalteromonas* is a novel symbiont with substantial evidence of phylosymbiosis across at least three marine invertebrate phyla (e.g., Nematoda, Mollusca, and Cnidaria). Patterns of symbiosis were consistent irrespective of geography (including in Antarctica), with FISH images from nematodes indicating that bacterial symbionts form biofilms in the mouth and esophagus. The evolutionary history of *Pseudoalteromonas* is marked by substantial host-switching and lifestyle transitions, and host-associated genomes suggest that these bacteria are facultative symbionts involved in nutritional mutualisms. In marine environments, we hypothesize that horizontally-acquired symbionts may have co-evolved with invertebrates, using host mucus as a physical niche and food source, while providing their animal hosts with Vitamin B, amino acids, and bioavailable carbon compounds in return.

## Introduction

Phylosymbiosis is commonly defined as “microbial community relationships that recapitulate the phylogeny of their host” (S. J. Lim & Bordenstein, 2020), but this strict definition is often expanded to include cophylogenetic signals (where related hosts interact with closely related bacterial taxa, often producing complex cophylogenetic topologies) and host-symbiont codiversfication (where two or more interacting species undergo reciprocal natural selection; (Dismukes et al., 2022)). Assessing phylosymbiosis in marine species is particularly challenging owing to the more “diffusible” nature of marine environments (dilution of bacterial cells and metabolites; (O’Brien et al., 2019; Shnit-Orland & Kushmaro, 2009)), the likely co-occurrence of specialist symbiont taxa alongside more generalist invertebrate microbiome assemblages (O’Brien et al., 2019; Shnit-Orland & Kushmaro, 2009), and the presumed horizontal transmission of many putative marine symbionts, which complicates efforts to identify clear genomic signatures of symbiosis and distinguish host-associated bacteria from free-living species, in contrast to the defined evolutionary constraints and genome reductions of vertically-transmitted symbionts (Boscaro et al., 2017; McCutcheon & Moran, 2011; Moran, 2007). In addition, invertebrate host species can flexibly acquire locally adapted symbiont strains (Breusing et al., 2022; Morrow et al., 2015) or consistently associate with the same symbiont lineage on a global scale (Neave, Rachmawati, et al., 2017), indicating that geographic and evolutionary patterns of symbiosis are highly variable and must be examined at the species level for both the host and microbe.

Does phylosymbiosis exist in marine invertebrate microbiomes? While there have been growing insights into the composition of marine invertebrate microbiomes using 16S rRNA marker gene surveys (Holt et al., 2023; Leasi et al., 2024; McCauley et al., 2023), our understanding of the broader evolutionary and ecological patterns that govern marine microbiome assembly remains poor. In the largest study carried out to date, Boscaro et al. (2022) found no evidence of phylosymbiosis across ∼1000 marine invertebrate microbiomes spanning 21 animal phyla, despite invertebrate microbiomes clustering separately overall from environmental microbial assemblages. In contrast, more targeted studies have found compelling evidence of phylosymbiosis in diverse animal taxa such as sponges (O’Brien et al., 2019, 2020), corals (O’Brien et al., 2019, 2020; Prioux et al., 2024), nemertean worms (Leasi et al., 2024), and brachyuran crabs (Tsang et al., 2024), even when using data from a single marker gene.

These conflicting patterns suggest that 16S rRNA surveys are inadequate for evaluating phylosymbiosis in marine environments, owing to the low resolution of marker gene datasets and an unfavorable signal-to-noise ratio (since rRNA datasets also capture signals from extracellular DNA, gut contents, and transient microbiome taxa alongside any putative symbiont taxa; (Bairoliya et al., 2022; T. J. Pereira et al., 2020)). Taken together, the above issues highlight that evaluating phylosymbiosis in marine invertebrates remains a major challenge.

There is a critical need for targeted genome-scale studies to investigate phylosymbiosis in marine invertebrates, incorporating expanded sampling of animal hosts as well as phylogenomic analyses of suspected bacterial/archaeal symbiont lineages.

Bacterial groups previously identified as “core microbiome taxa” in 16S rRNA metabarcoding studies represent an ideal starting point for deeper investigations of phylosymbiosis in marine invertebrates (O’Brien et al., 2019). Here, we use novel cultured isolates and pangenomic analyses to investigate *Pseudoalteromonas* as a ubiquitous bacterial genus that may represent a novel and previously unrecognized symbiont of marine invertebrates, with high potential for host-microbe coevolution. *Pseudoalteromonas* is consistently recovered as an abundant (and often, statistically significant) taxon in diverse invertebrate microbiomes, including marine nematodes (Bellec et al., 2019; Moens et al., 2005), gelatinous zooplankton (McCauley et al., 2023; Ohdera et al., 2023; T. J. Pereira et al., 2023), corals (Delgadillo-Ordoñez et al., 2024; Osman et al., 2020), sponges (Williams et al., 2024), shrimp (Shan et al., 2025), molluscs (Zhu et al., 2024), and even some protists, including bloom-forming dinoflagellates (Seibold et al., 2001). *Pseudoalteromonas* bacteria are ubiquitous across marine ecosystems, comprising free-living and particle-associated taxa (isolated from surface ocean waters down to deep-sea sediments; (Salazar et al., 2015; Wietz et al., 2010)) as well as host-associated species, with the first cultured isolate originally obtained from a marine alga in 1957 (Yaphe, 1957). Subsequent host-associated isolates have been acquired from pelagic and benthic marine invertebrates, including both sessile and mobile species (Atencio et al., 2018) as well as bony fish (Bell et al., 2024). Similar to other well-characterized symbionts (Adnani et al., 2017), *Pseudoalteromonas* species are characterized by a high degree of bioactivity, producing a variety of secondary metabolites and antibiotics that inhibit the growth of other bacterial taxa (Atencio et al., 2018; Bosi et al., 2017). *Pseudoalteromonas* are often found as mucus-associated taxa in corals (O’Brien et al., 2019; Shnit-Orland & Kushmaro, 2009) and have even been observed as the sole bacteria able to colonize marine nematode mucus tracts (suggesting exploitation of mucus as a unique physical niche; (Moens et al., 2005)).

*Pseudoalteromonas* is also known to influence the development and fitness of several marine invertebrate species: stimulating immune response, enhancing resistance to pathogenic infections, and increasing the digestive enzyme activity of *Apostichopus japonicus* sea cucumber hosts (Ma et al., 2014), and containing genomic pathways that induce larval settlement and metamorphosis of several marine invertebrate species including the upside-down jellyfish *Cassiopea xamachana* (Ohdera et al., 2023), the tubeworm *Hydroides elegans* (Ericson et al., 2019; Malter et al., 2025), and the mussel *Mytilus coruscus* (Peng et al., 2020). Over evolutionary timescales, *Pseudoalteromonas* bacteria may have established species-specific symbiotic relationships with diverse marine invertebrates, deploying robust metabolic toolkits to establish a host-associated niche and providing distinct benefits to marine invertebrates.

## Results

### An expanded pangenomic view of the ubiquitous bacterial genus Pseudoalteromonas

To characterize the genomic repertoire and functional diversity of the genus *Pseudoalteromonas*, we constructed a pangenome using 234 publicly available genomes and two novel nematode-associated *Pseudoalteromonas undina* strains isolated as part of this study (**Supplementary Table 1**), the latter of which we obtained by adapting high-throughput culturing approaches originally developed for studies of pelagic marine bacteria (De Santiago et al., 2025; Henson et al., 2020). Our study represents a drastic expansion of two previous *Pseudoalteromonas* pangenomic analyses that used fewer than 38 genomes to characterize the evolutionary history of this ubiquitous bacterial clade (Bosi et al., 2017; Sonnenberg & Haugen, 2021). Our final pangenome consisted of 91 free-living and 145 host-associated *Pseudoalteromonas* isolated from various marine environments. The host-associated strains were previously isolated from nine marine invertebrate Phyla (Arthropoda, Annelida, Chordata, Cnidaria, Ctenophora, Echinodermata, Nematoda, Mollusca, and Porifera; **Fig. 1A**). The most common hosts were Cnidaria (78 isolates), followed by Mollusca (19 isolates), and Porifera (18 isolates). The phylum Annelida (1 isolate), Nematoda (2 isolates, both obtained in this study), and Chordata (3 isolates) were the least common animal hosts in our dataset.

**Fig. 1:**
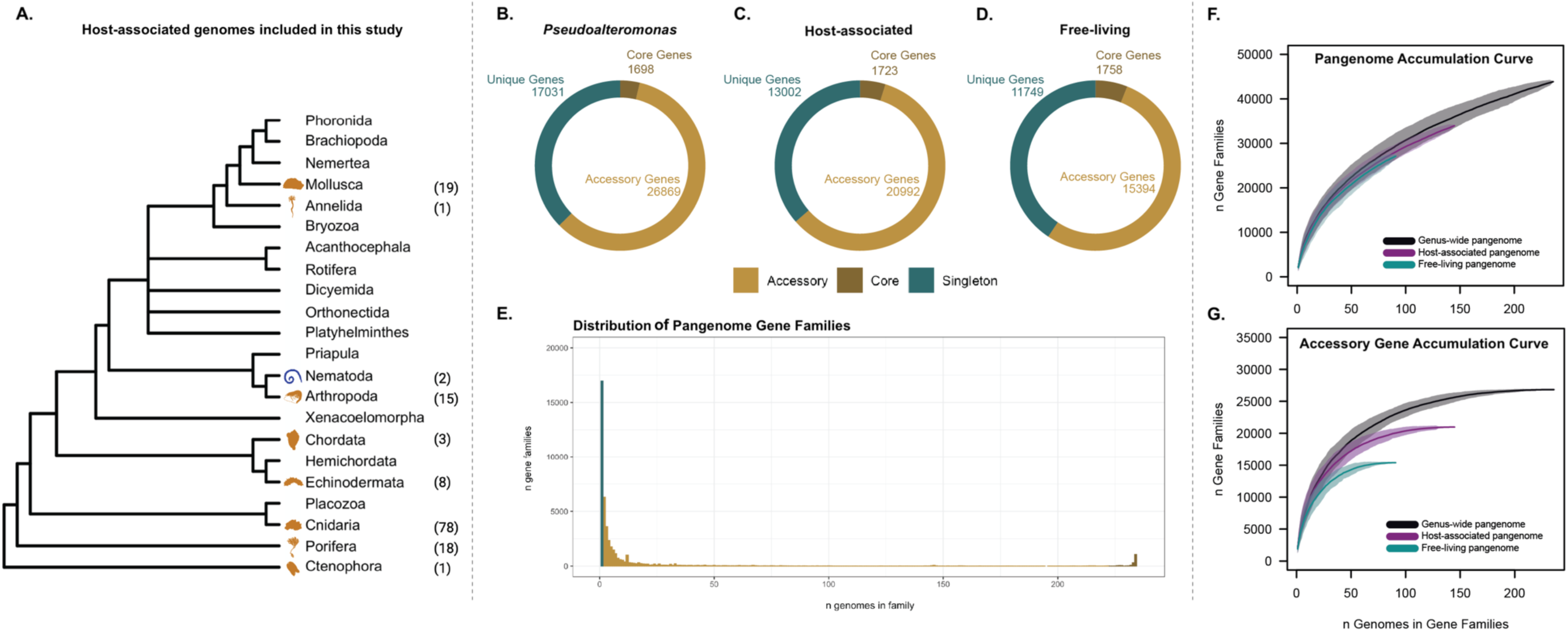
*Pseudoalteromonas* has an open pangenome that is driven by the accumulation of unique genes. **A**: The phylogenetic tree of marine invertebrates was adapted from Boscaro et al. (2022). The Blue PhyloPics (e.g., Nematoda) indicates host-associated *Pseudoalteromonas* that were isolated from this study, and gold PhyloPics (e.g., Annelida, Arthropoda, Chordata, Cnidaria, Ctenophora, Echinodermata, Mollusca, and Porifera) indicate the publicly available host-associated *Pseudoalteromonas* genomes included in this study. Parenthetical numbers represent the number strains isolated from each phylum. **B-D**: The number of core genes (brown; 95% of genomes), accessory genes (gold; present <95% of genomes), and unique genes (green) found in B) all *Pseudoalteromonas*, C) Host-associated, and D) Free-living pangenomes. E) The distribution of genes in the *Pseudoalteromonas* pangenome. F) Accumulation curves of the three pangenomes indicate that *Pseudoalteromonas* has a large pangenome; however, G) accumulation curves of the accessory genes reach an asymptote, indicating that the pangenome openness is being driven by unique genes.

The size and distribution of the *Pseudoalteromonas* pangenome were calculated using three parallel approaches: 1) the entire genus, 2) free-living genomes only, and 3) host-associated genomes only. Following a standard approach used to characterize bacterial pangenomes (Barh et al., 2020), gene families were divided into three groups according to the frequency at which they appear in the pangenome (**Fig. 1B-D**): core genes (present in >95% of the genomes), accessory genes (present in <95% of the genomes but not unique to a single genome), or unique genes (found only in one genome). Our results showed that the *Pseudoalteromonas* pangenome could be classified as an open pangenome, characterized by a small set of core genes (3.72%; 1,698 genes) and a large number of accessory (58.93%; 26,869) and unique genes (37.35%; 17,031). When partitioned by lifestyle (**Fig. 1C-D**), the host-associated pangenome contained 5,598 more accessory genes (20,992) compared to the free-living pangenome (15,394; p-value < 2.2e-16). Accumulation curves of the entire pangenome, which include the core, accessory, and unique gene families, did not asymptote, indicating that *Pseudoalteromonas* has an open pangenome and that our expanded pangenomic analysis does not yet capture the true biological diversity of this genus (**Fig. 1F**). However, accumulation curves of the accessory gene families for the free-living, host-associated, and genus-level pangenomes reached an asymptote (**Fig. 1G**), indicating that the openness of the pangenome was being driven mainly through the accumulation of unique genes. Our expanded pangenome analysis is consistent with previous findings that identified a remarkable genomic heterogeneity within this bacterial genus, even for closely related *Pseudoalteromonas* strains (Bosi et al., 2017). These results indicate that *Pseudoalteromonas* lineages maintain a high degree of genomic flexibility, which may facilitate rapid exploitation of diverse ecological niches in the marine environment.

### Pseudoalteromonas exhibits distinct phylogroups and frequent lifestyle transitions

To unravel the evolutionary dynamics and population structure of free-living and host-associated *Pseudoalteromonas*, core genes (1,698) were used to construct a midrooted maximum likelihood phylogenetic tree and genomes were delineated into phylogroups using average nucleotide identity (ANI). We identified 57 distinct phylogroups, substantially increasing the known diversity of *Pseudoalteromonas* beyond the 34 species currently described in the literature (**Fig. 2A and Extended Data Fig. 1**). Across the phylogenetic tree, phylogroup membership and clade branching patterns indicated frequent transitions between free-living and host-associated lifestyles as well as substantial host-switching between invertebrate taxa (**Fig. 2A**), indicative of a flexible genomic repertoire in *Pseudoalteromonas* that facilitates association with diverse hosts and habitats. These evolutionary patterns suggest a “nomadic” lifestyle seen in other bacterial groups (where bacteria maintain a “universal” set of genes that allow them to thrive in a variety of environments; (Martino et al., 2016)), and phylogenetic patterns indicate an absence of clade-specific environmental adaptation. Of the 57 phylogroups, 35.09% (20 phylogroups) comprised a mixture of host-associated and free-living isolates. The majority of the phylogroups (64.91%) were lifestyle-specific, containing only free-living or only host-associated taxa. Of the lifestyle-specific phylogroups, 25 comprised only host-associated genomes, versus fewer than 12 phylogroups with entirely free-living strains. However, the majority of these lifestyle-specific phylogroups (37) contained less than three genomic representatives, suggesting poor taxon sampling and the need for additional sampling efforts to confirm the veracity of these patterns. Overall, the *Pseudoalteromonas* phylogeny was divided into three major clades: a group of closely related bacteria with shorter branch lengths (nonpigmented) and two smaller clades with longer branches and greater evolutionary divergence (pigmented; **Fig. 2A**). The structure and topologies of these two major clades reflects previously described phylogenetic patterns for pigmented and non-pigmented clades of *Pseudoalteromonas* species (Bosi et al., 2017).

**Fig. 2:**
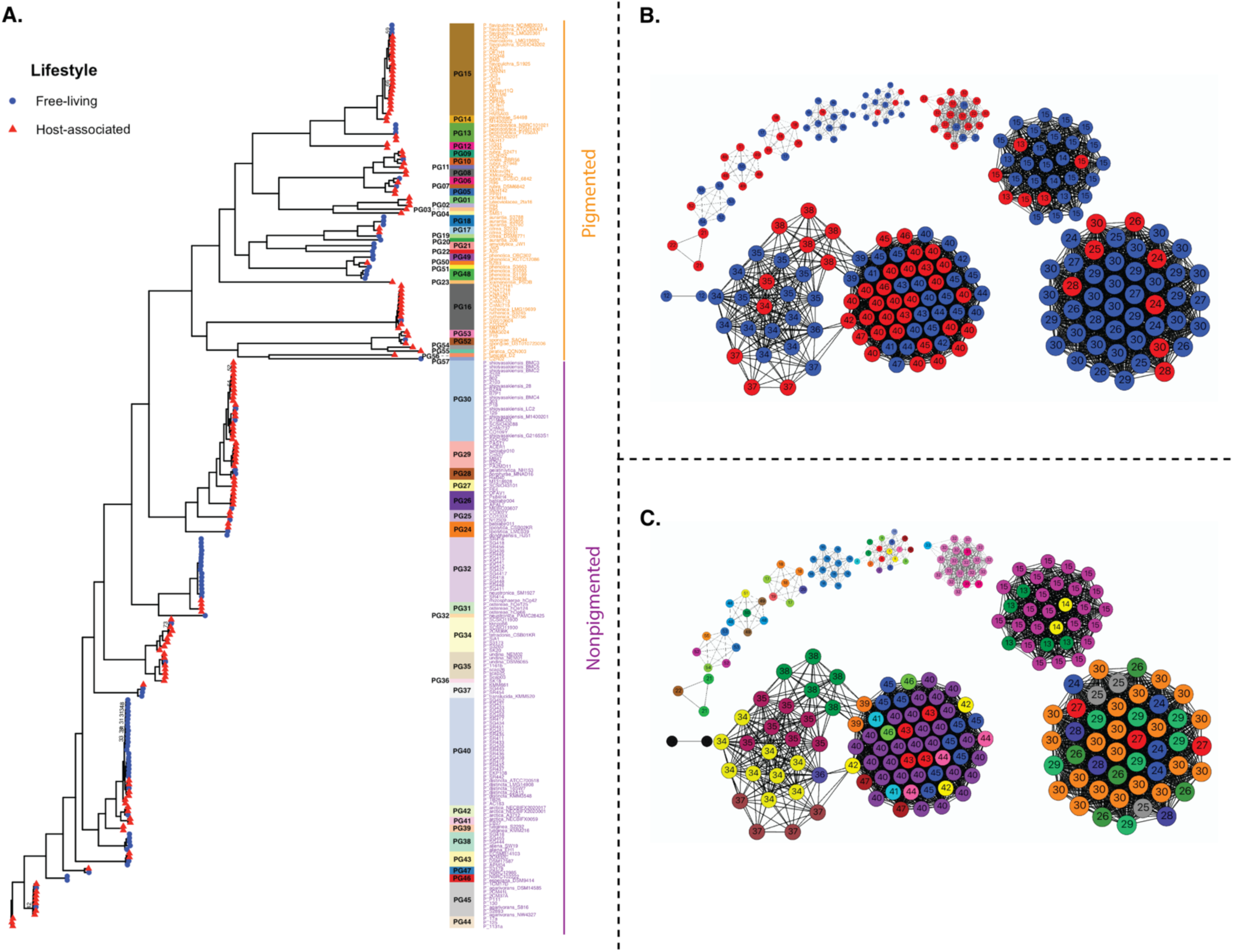
The accessory pangenome of *Pseudoalteromonas* clusters according to phylogroups, rather than bacterial lifestyle. A) An unrooted phylogenetic tree constructed using the 1,689 core genes and 10,000 bootstraps. Tips with red triangles indicate host-associated genomes, while blue circles indicate free-living genomes. The labels indicate whether the clades are pigmented (orange) or non-pigmented (purple), according to a previous pangenomic study (Bosi et al., 2017). B) Genome-genome network analysis using most of the accessory genome (removing accessory genes present in <5% of the genomes). Red and blue circles indicate host-associated and free-living genomes, respectively. C) The genome-genome network analysis with samples labeled according to phylogroup. The colors are independent of panel A.

To assess whether the distribution of accessory gene families corresponded to lifestyle (host-associated vs. free-living) or evolutionary history (phylogroups), we conducted a genome-genome network analysis using accessory genes found in at least 5% of the genomes (**Fig. 2B-C**). The isolates were not grouped by lifestyle but were instead structured by closely related phylogroups (**FIG. 2B-C**). The genome size (Wilcoxon p-value = 0.87) and number of genes (Wilcoxon p-value = 0.67) between host-associated and free-living genomes were not significantly different (**Fig. 3A-B**). However, the host-associated genomes exhibited significantly higher coding density (Wilcoxon p-value < 0.01) and a nucleotide bias (higher GC Content; Wilcoxon p-value < 0.01) compared to the free-living genomes (**Fig. 3C-D**). Such evolutionary “jumps” in GC content – including both increases and decreases in GC content – have been increasingly documented in bacterial lineages with host-associated lifestyles (Mahajan & Agashe, 2022), although the mechanisms explaining such GC jumps are not well understood.

**Fig. 3:**
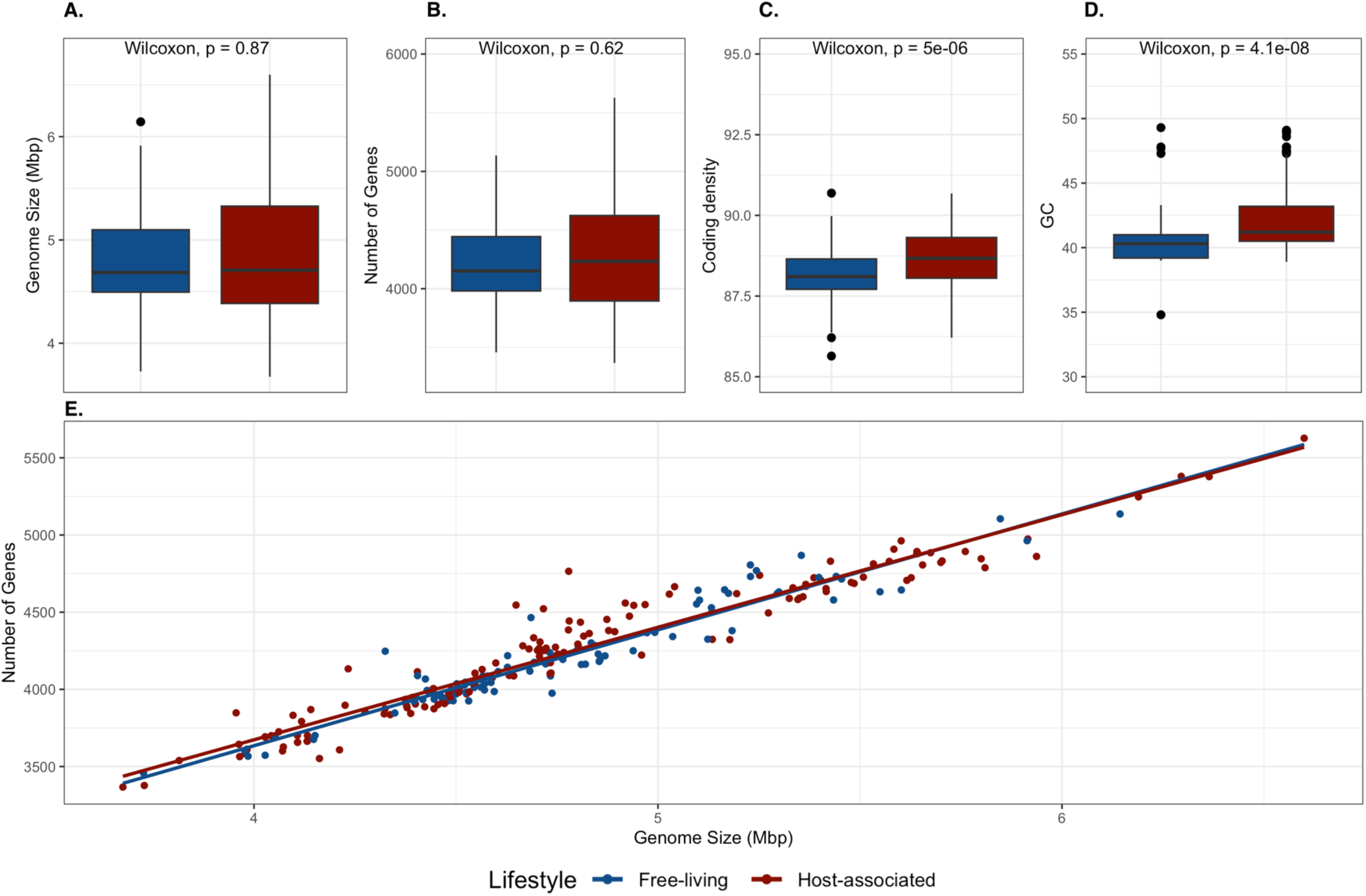
Host-associated lineages exhibit significantly higher coding density and GC content compared to free-living strains. Differences between four different genomic characteristics, including A) Genome Size (bp), B) Total Number of Genes, C) Coding Density, and D) GC Content (%) between host-associated and free-living *Pseudoalteromonas* genomes. Differences coding density and GC content were significant based on Wilcoxon signed-rank tests.

### Evidence of phylosymbiosis across diverse marine invertebrates

To test for phylosymbiosis between marine invertebrates and *Pseudoalteromonas,* we constructed seven host and bacterial phylogenetic trees: one tree spanning all hosts in the Kingdom Animalia (to evaluate broad patterns of phylosymbiosis, analogous to Boscaro et al. (2022)), four phylum-specific trees for invertebrate groups representing the most well-sampled animal phyla in our dataset (Mollusca, Cnidaria, Porifera, and Nematoda), and trees for class Bivalvia (Mollusca) and family Faviidae (Cnidaria) representing the two host phylogenies with the most extensive taxon sampling at lower taxonomic levels.

Using genome-scale data, we found consistent evidence of phylosymbiosis at most of the taxonomic levels examined (**Fig. 4, Extended Data Fig. 2, and Table 1**), despite *Pseudoalteromonas* genomes being collected across different temporal and spatial scales. At the level of Kingdom Animalia, we found evidence of low cophylogenetic signal, but not of phylogenetic congruence (Generalized RF = 0.78; PACo p-value < 0.001; PACo R^2^ = 0.07; **Extended Data Fig. 2**). This low (but statistically significant) cophylogenetic signal across all animal phyla is likely due to substantial host-switching by *Pseudoalteromonas* lineages among phylogenetic groups as phylogenetic congruences and cophylogenetic signals reflect two different evolutionary processes (**Extended Data Fig. 2**). Phylogenetic congruence measures whether the host’s evolutionary history is dependent on the symbiont (or vice-versa); however, cophylogenetic signals can emerge if the host (or symbiont) evolutionary history is shaped by conserved traits or biogeography (Perez-Lamarque & Morlon, 2024). Statistical support for phylosymbiosis becomes stronger when animal hosts are evaluated at the phylum, class, and family levels. We found evidence of cophylogeny and weak phylogenetic congruence between *Pseudoalteromonas* and Mollusca (Generalized RF = 0.60; PACo p-value = 0.012; PACo R^2^ = 0.60; **Fig. 3A and Table 1**). In Mollusca, two bacterial lineages belonging to phylogroup PG15 interacted with two cephalopod hosts, while PG30 showed evidence of host-switching from Crassostrea (true oysters) to Haliotis (abalone sea snails). Within class Bivalvia (Mollusca), we recovered high cophylogenetic signal and weak phylogenetic congruence with *Pseudoalteromonas* (Generalized RF = 0.75; PACo p-value = 0.025; PACo R^2^ = 0.41; **Fig. 3B and Table 1**), where five *Pseudoalteromonas* lineages (spanning three phylogroups) interacted with Crassostrea and exhibited host-switching with two other bivalve groups, Mytilidae (mussels) and *Ostrea* spp. (edible oysters). There was low cophylogenetic signal but no phylogenetic congruence between *Pseudoalteromonas* and the phylum Cnidaria (Generalized RF = 0.85; PACo p-value = 0.011; PACo R^2^ = 0.11; **Fig. 3C and Table 1**), but high cophylogenetic signal and weak phylogenetic congruence for stony corals within family Faviidae (Generalized RF = 0.73; PACo p-value = 0.025; PACo R^2^ = 0.41; **Fig. 3D and Table 1**). Within the Faviidae, three *Pseudoalteromonas* lineages (PG15, PG9, and PG50) interacted closely with *Diploria* spp. (grooved brain corals), and five *Pseudoalteromonas* lineages (PG16, PG24, PG26, PG29, PG30) interacted with *Colpophyllia* spp. (boulder brain corals). Additionally, the two *Pseudoalteromonas* lineages associated with *Pseudodiploria* spp. stony corals demonstrated host switching. Similar to other invertebrate phyla, we saw evidence of high cophylogenetic signal between *Pseudoalteromonas* and the phylum Nematoda (PACo p-value < 0.001; PACo R^2^ = 0.20; **Fig. 5 and Table 1**) when assessing host/bacterial phylogenetic trees. Only one analysis showed no evidence of cophylogenetic signal or phylogenetic congruence: that between *Pseudoalteromonas* and the phylum Porifera (Generalized RF = 0.94; PACo p-value = 0.470; PACo R^2^ = 0.14; **Table 1**). Evidence of phylosymbiosis became stronger (higher cophylogenetic signal and the emergence of statistically significant phylogenetic congruence) when lower-level taxonomic groups were assessed (class Bivalvia and family Faviidae).

**Fig. 4:**
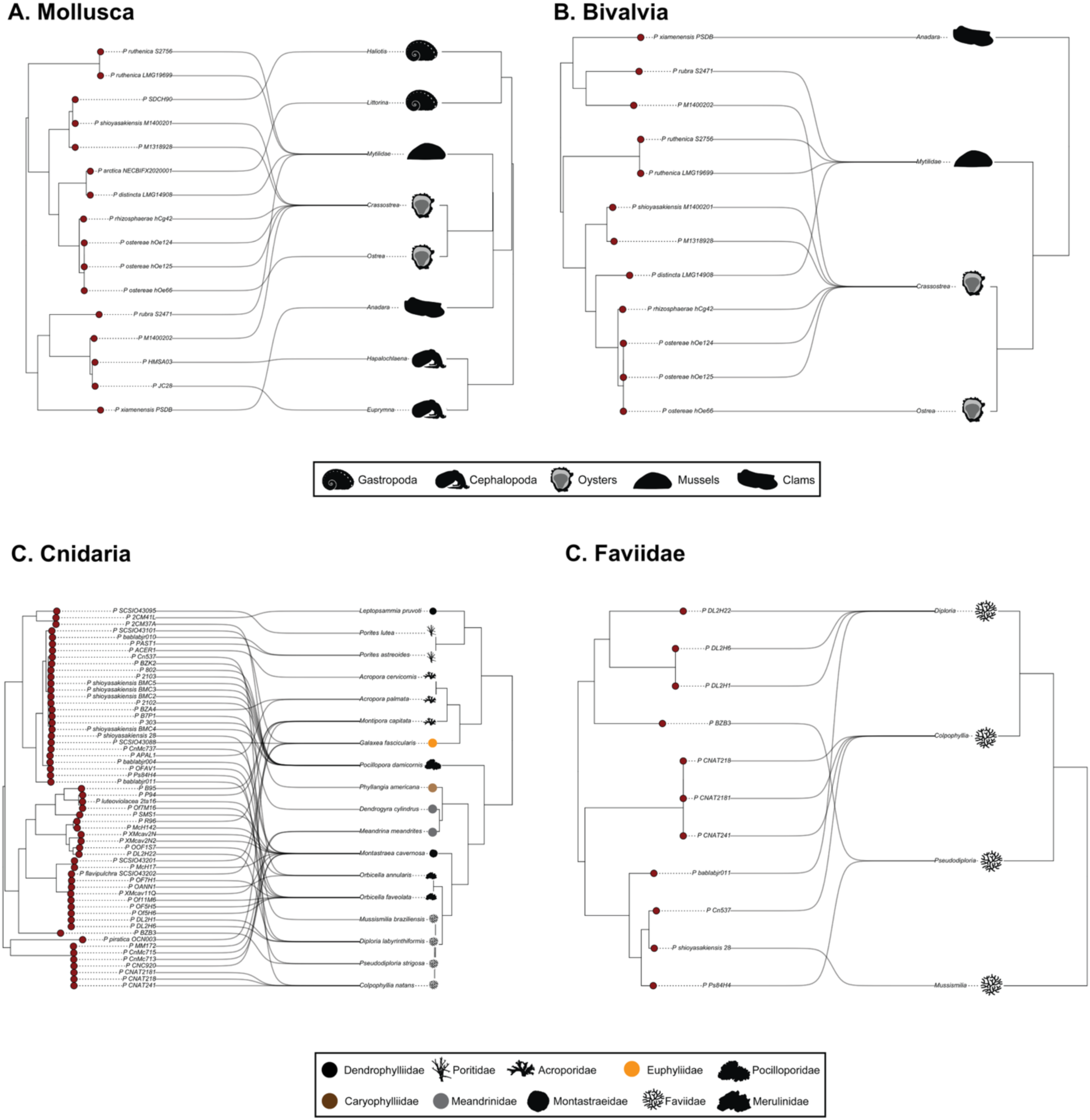
Evidence of cophylogeny and phylogenetic congruence between *Pseudoalteromonas* and marine invertebrate taxa. Phylogenetic trees used to test for cophylogeny among A) Mollusca, B) Bivalvia (Mollusca), C) Cnidaria, and D) Faviidae (Cnidaria). Cophylogeny signal was tested using PACo, and phylogenetic congruence was tested using the Generalized Robinson-Foulds metric (**Table 1**).

**Fig. 5:**
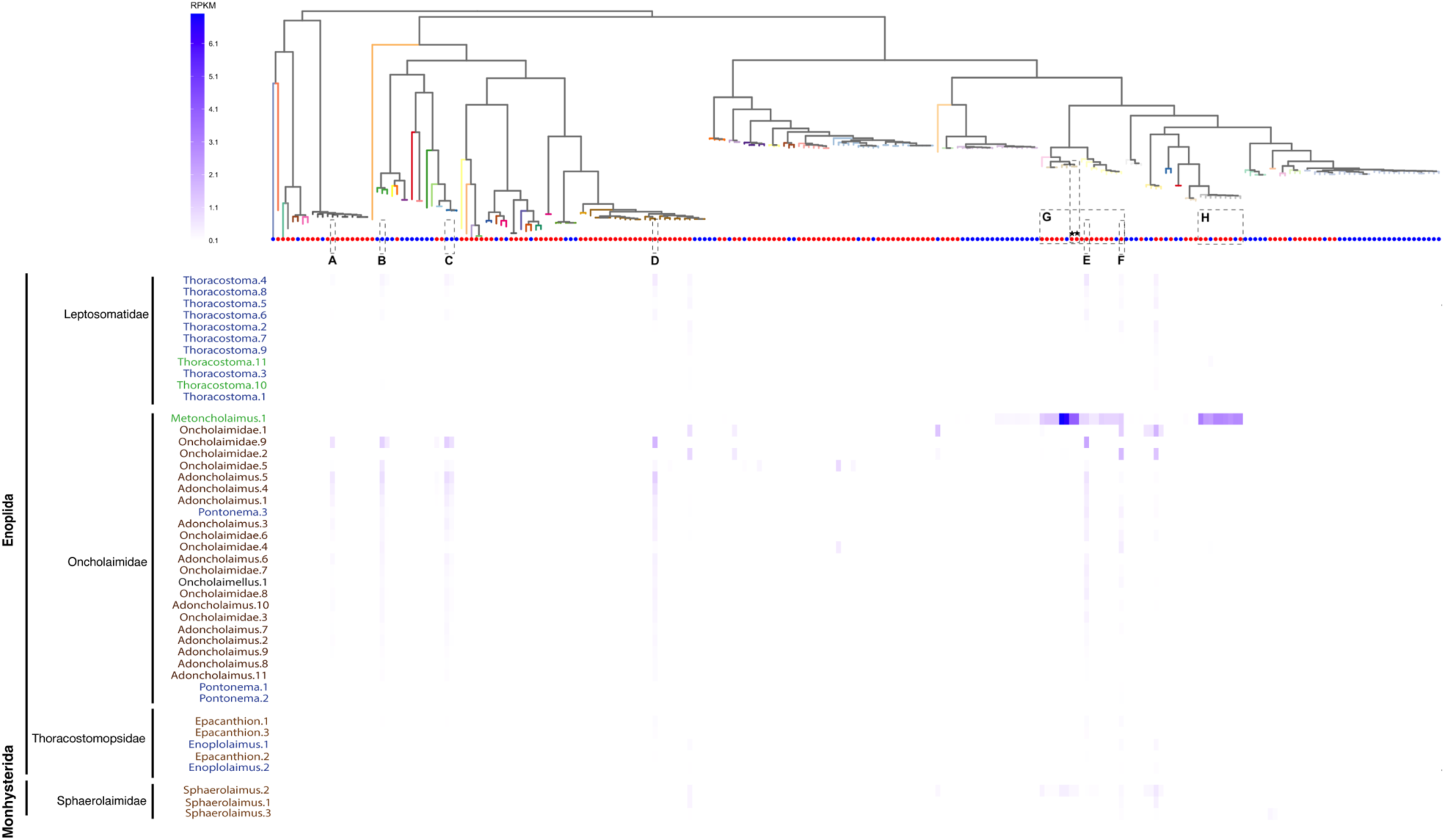
Read-mapping approaches recover consistent signals from novel Pseudoalteromonas symbionts irrespective of geographic region. Heatmap depicts metagenome reads from single-worm sequencing that map to *Pseudoalteromonas* genomes. The *Pseudoalteromonas* bacterial phylogeny is displayed horizontally across the top, with branch coloring indicating phylogroup membership and circles indicating lineages with host-associated (red) or free-living (blue) lifestyles. Black stars indicate the two isolates obtained as part of this study. Nematode metagenomes are labelled on the left-hand side, arranged by taxonomic hierarchy (bold indicates order, plain black text indicates family). Marine nematode samples are identified down to family or genus of each individually sequenced worm and colored according to the geographic location where each specimen was obtained: : Tybee Island, GA (Brown), Bodega Bay, CA (Green), Dolphin’s Reef, FL (Gray), and Antarctica (Blue).

**Table 1.** Statistical support for phylosymbiosis in marine invertebrates, indicated by cophylogenetic signals and phylogenetic congruence of *Pseudoalteromonas*. Results from the PACo analysis and the Generalized Robinson-Foulds metric indicate whether there is a cophylogenetic signal (PACo p-value < 0.05) and evidence of phylogenetic congruence across marine invertebrate phyla. A generalized RF of 0 indicates perfect congruence between host-associated and bacterial phylogenies.

| Host | PACo<br>P-value | PACo<br>m<sup>2</sup> | PACo<br>R<sup>2</sup> (1 - m<sup>2</sup>) | Generalized<br>Robinson-Foulds | Cophylogenetic<br>Signal | Phylogenetic<br>Congruence |
| --- | --- | --- | --- | --- | --- | --- |
| Animalia | 0.00 | 0.93 | 0.07 | 0.85 | Low | None |
| Mollusca | 0.01 | 0.60 | 0.40 | 0.60 | High | Weak |
| Cnidaria | 0.01 | 0.89 | 0.11 | 0.85 | Low | None |
| Porifera | 0.47 | 0.86 | 0.14 | 0.93 | None | None |
| Nematoda | 0.00 | 0.80 | 0.20 | NA | High | None |
| Bivalvia<br>(Mollusca) | 0.01 | 0.65 | 0.34 | 0.75 | High | Weak |
| Faviidae<br>(Cnidaria) | 0.03 | 0.59 | 0.41 | 0.73 | High | Weak |

However, analysis at lower taxonomic levels only included four-member host trees indicating that higher taxon sampling is needed to confirm phylosymbiosis (although 4-member trees have previously been sufficient to assess clade-specific phylosymbiosis; (Brooks et al., 2016)). High cophylogenetic signal was also obtained at the phylum level (in Mollusca and Nematoda) when taxon sampling was deep enough for statistical support to emerge. These data suggest that bacterial genome data, in addition to extensive host taxon sampling, are imperative for detecting signals of phylosymbiosis in marine invertebrates. The frequent lifestyle transitions of *Pseudoalteromonas* lineages and apparent host-switching seen across marine invertebrates can further obscure signals of phylosymbiosis at the kingdom/phylum level, particularly in previous studies that utilize only low-resolution 16S rRNA data (Boscaro et al., 2022).

To further assess phylosymbiosis between *Pseudoalteromonas* and representatives from the phylum Nematoda, we obtained nematode-associated microbiomes via single-worm sequencing from eight marine nematode genera (representing two orders and four families). Samples were collected from four distinct environments (Antarctic continental shelf, intertidal sites on the Georgia coast, subtidal coral sands in the Florida Keys, and seagrass habitats in Bodega Bay, California; **Supplementary Table 2**) and bacterial abundances were quantified as Reads Per Kilobase per Million mapped reads (RPKM). Single-nematode read-mapping indicated that the same *Pseudoalteromonas* strains were consistently associated with nematode hosts irrespective of geography: Antarctic nematode microbiomes contained the same *Pseudoalteromonas* lineages as nematode specimens obtained from the Georgia, California, and Florida coastlines (Fig 5A-F). Notably, we often detected five or more distinct *Pseudoalteromonas* lineages associated with individual marine nematodes (spanning both pigmented and non-pigmented bacterial clades; **Fig. 5A-E**), suggesting the existence of complex symbioses involving multiple congeneric bacterial species. *Pseudoalteromonas* read-mapping signals were strongest within the nematode family Oncholaimidae (Mean RPKM: 5.94) and Sphaerolaimidae (Mean RPKM: 3.44), consistent but weaker in family Leptosomatidae (Mean RPKM: 1.29), and sporadic or absent in Thoracostomopsidae (Mean RPKM: 0.97; **Fig. 5B-F**), further suggesting that associations with *Pseudoalteromonas* may be particularly important for the ecology and life history of a specific subset of marine nematode taxa, and these bacteria may have co-evolved with specific evolutionary lineages of nematodes (e.g., family Oncholaimidae). Of the *Pseudoalteromonas* lineages detected with read-mapping in family oncholaimid nematode specimens, three were previously characterized as free-living *Pseudoalteromonas* strains, while four were previously isolated from invertebrate hosts (three from Arthropoda and one from Cnidaria; **Fig. 5A-F**), indicating that our overall knowledge of *Pseudoalteromonas* habitats and host associations is incomplete. In addition, metagenomic reads from one oncholaimid nematode genus (*Metoncholaimus* spp.) mapped to all the strains in several *Pseudoalteromonas* phylogroups (PG45, PG43, PG34, PG35, and PG36), which include free-living and host-associated strains previously isolated from various invertebrate taxa (Arthropoda, Chordata, Cnidaria, Echinodermata, Mollusca and Porifera), including the two new nematode-associated *Pseudoalteromonas* isolates obtained in this study (**Fig. 5G-H**). A single sample from the family Sphaerolaimidae (Sphaerolaimus.2) mapped to some of the same phylogroups recovered in the *Metoncholaimus* spp. nematode (PG43, PG35, PG36), including the two nematode-associated strains obtained in the present study (**Fig. 5**). These patterns suggest that marine nematodes may associate with a phylogenetically diverse set of *Pseudoalteromonas* lineages spanning many phylogroups (as seen for most marine nematodes in family Oncholaimidae and Leptosomatidae) or alternatively harbor many closely-related *Pseudoalteromonas* lineages restricted to a small number of bacterial phylogroups (as seen in the oncholaimid genus *Metoncholaimus*). Most importantly, patterns of nematode-associated *Pseudoalteromonas* were consistent within family-level clades and maintained across vast geographic distances, suggesting that these novel symbioses have ancient evolutionary origins.

### Functional pathways suggest nutritional mutualism in host-associated lineages

We used a panGWAS (Roder et al., 2024) approach to identify genes and functions that were more common in host-associated genomes. Due to substantial differences in evolutionary rates between the pigmented and non-pigmented *Pseudoalteromonas* genomes, we analyzed these clades separately to identify genes that may underpin host-associated lifestyles, as recommended by a prior pangenomic study (Bosi et al., 2017). Lifestyle transitions towards symbiosis appear to emphasize genes and pathways related to the breakdown of organic compounds (peptides, cellulose, chitin, and other carbohydrates), production and transport of antimicrobial and antitoxin compounds, biofilm formation, production of essential vitamins, and enhanced transport of Vitamin B12.

Of the 26,869 pigmented accessory genes, only six were more abundant in the host-associated pigmented strains (three of which were hypothetical proteins; **Extended Data Fig. 3 and Supplementary Table 3**). The three annotated genes included an inner membrane transport protein YhdP (panGWAS odds-ratio = 8.75; p-value < 0.001) that maintains outer membrane permeability, an acetoin utilization protein (panGWAS odds-ratio = 22.15; p-value < 0.0001), and a HTH-type transcriptional regulator DmlR (panGWAS odds-ratio = 8.09; p-value = 0.001) that regulates aerobic growth on D-malate (**Supplementary Table 3**). The three AI-annotated hypothetical proteins (see methods) were predicted to be a periplasmic substrate-binding protein (panGWAS odds-ratio = 12.00; p-value = 0.0001), a complex carbohydrate transporter (panGWAS odds-ratio = 8.70; p-value = 0.0001), and a gene involved in bacteriocin (antimicrobial peptide) production or immunity (panGWAS odds-ratio = 8.413; p-value = 0.0001).

In the non-pigmented genomes, 111 genes were more abundant in the host-associated strains, with 70 classified as hypothetical proteins (**Supplementary Table 3**). The annotated genes included three additional HTH-type transcriptional regulators (panGWAS odds-ratio > 3.7; p-value < 0.001), the complete pathway for three hydrogen cyanide synthase subunits (panGWAS odds-ratio > 5.03; p-value = 0.0001), four Vitamin B12 transporters (panGWAS odds-ratio = 4.31-5.67; p-value < 0.0001), and multidrug transporters (panGWAS odds-ratio = 6.67; p-value < 0.000). Additionally, two glycoside hydrolases, a chitinase (panGWAS odds-ratio = 5.58; p-value < 0.0001) and a chitodextrinase (panGWAS odds-ratio = 4.32; p-value < 0.0001), were also more common in the nonpigmented host-associated genomes. Of the 70 hypothetical proteins, five were predicted to play a role in toxin/antimicrobial systems (panGWAS odds-ratio = 4.32; p-value < 0.0001), and three genes were predicted to play a role in antibiotic resistance (panGWAS odds-ratio = 5.27 - 5.57; p-value < 0.0001). Additionally, two genes were predicted to be involved in amino acid transport (panGWAS odds-ratio = 4.89 - 5.56; p-value < 0.0002) and two were predicted by AI gene annotators to contribute to biofilm formation or cellular adhesions (panGWAS odds-ratio = 5.15 - 6.21; p-value < 0.0001). A single gene was predicted to play a role in phospholipid biosynthesis (panGWAS odds-ratio = 6.04; p-value = 0.0002). Genes involved in specialized nutrient and amino acid transport systems, fatty acid and lipid biosynthesis, and membrane-associated proteins are typically more common in bacteria that have a symbiotic lifestyle (Villada et al., 2025). Gene pathways also suggest that host-associated *Pseudoalteromonas* lineages may be able to utilize alternative carbon sources, such as malate, under aerobic conditions (Lukas et al., 2010; Xiong et al., 2019). Additionally, antimicrobial compounds and toxin/antitoxin systems are common in marine invertebrate microbiomes and may help host organisms resist pathogenic infections (Chen et al., 2021; L. B. Pereira et al., 2017).

We next used METABOLIC (Zhou et al., 2022) and antiSMASH (Blin et al., 2023) to predict metabolic pathways, biogeochemical functions, and secondary metabolites across the *Pseudoalteromonas* genomes in our dataset, to identify functions and genes that may facilitate host-microbe interactions (**Fig. 6 and Supplementary Table 4**). Several functions that facilitate bacteria-eukaryote interactions were commonly found across the entire pangenome (Douglas, 2017; Motta et al., 2024; Sultana et al., 2023). For example, all the *Pseudoalteromonas* genomes in our dataset are predicted to synthesize Vitamins B1 (Thiamin) and B2 (Riboflavin). Most bacterial lineages, with the exception of two genomes, are predicted to synthesize Vitamin B7 (Biotin), and approximately 60% of the genus can synthesize Vitamin B12 (cobalamin).

**Fig. 6:**
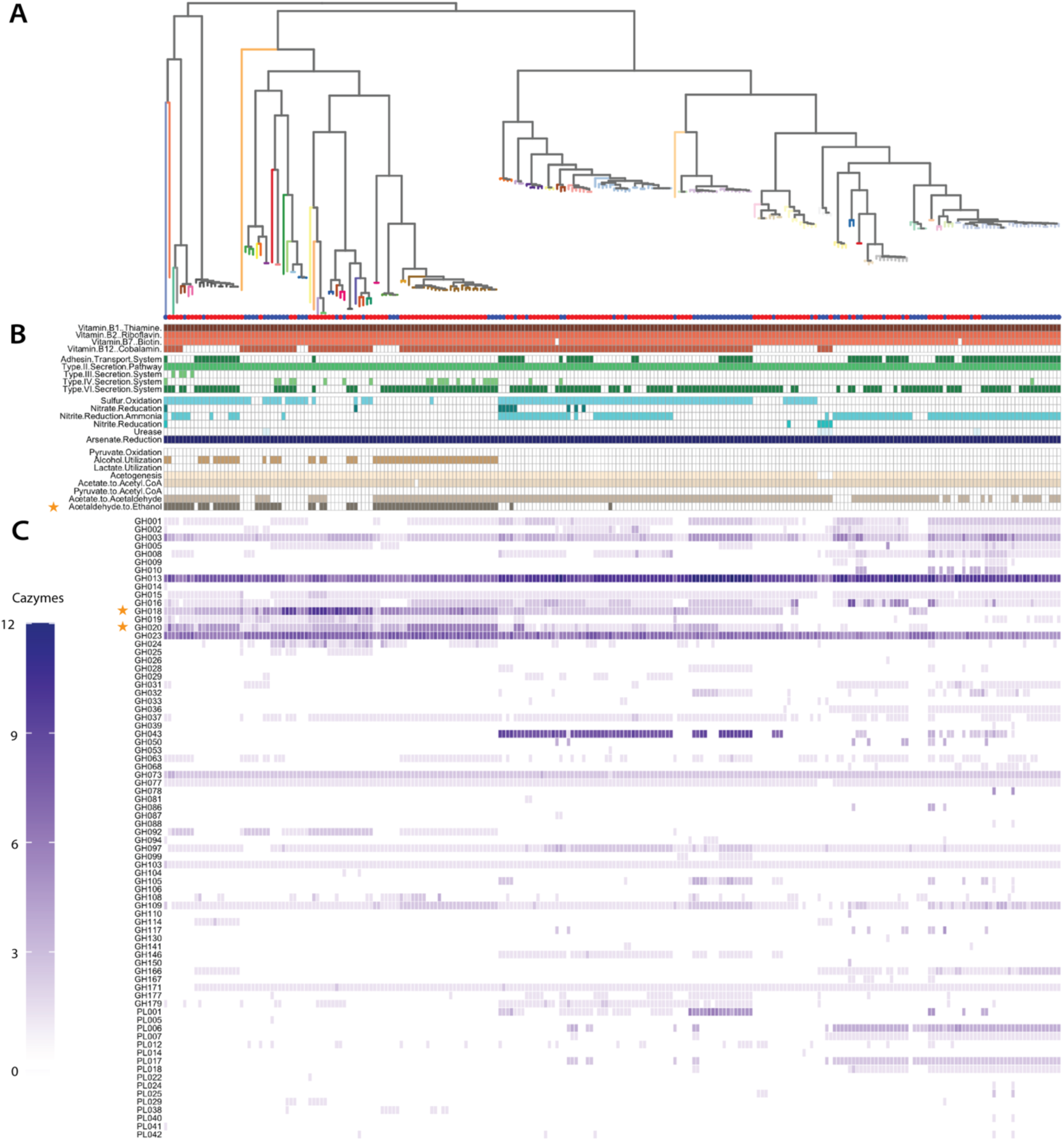
Host-associated *Pseudoalteromonas* lineages are enriched for genes and pathways related to B vitamin production, antimicrobial/antitoxin compounds, breakdown of organic molecules, and host-bacterial interactions. A) Midrooted phylogenetic tree of the *Pseudoalteromonas* pangenome. The tips indicated whether the isolate was a free-living (blue circle) or host-associated (red circle) strain, and the branch color indicates the phylogroup. B) A summary of the metabolic and biogeochemical traits, and C) the heatmap of the abundance of CAZymes predicted by METABOLIC (Zhou et al., 2022). Orange stars indicate functions that are enriched in host-associated pigmented genomes.

Additionally, about 70% of *Pseudoalteromonas* taxa (165 genomes) are predicted to contain a type VI secretion system, which is typically found in ∼52% of *Gammaproteobacteria* but is absent in the majority of bacterial phyla (Abby et al., 2016). Nearly half (∼48%) were predicted to perform sulfur oxidation via sulfur dioxygenases, an essential step in sulfide detoxification.

Some functions were conserved across the pangenome, including arsenate reduction, acetogenesis, and type II secretion (**Fig. 6**).

Only one endopeptidase (C56), was more common in host-associated genomes within the pigmented clade (panGWAS odds-ratio = 9.80; p-value = 0.002). In the non-pigmented clade, three functions were more abundant in the host-associated genomes: cellulose degradation (panGWAS odds-ratio = 7.23; p-value < 0.001), chitin degradation (panGWAS odds-ratio = 7.23; p-value < 0.001), and ethanol fermentation (panGWAS odds-ratio = 3.30; p-value = 0.003; (**Supplementary Table 4**)). Three peptidases (S09B, S01A, and C56; panGWAS odds-ratio = 3.30 - 5.60; p-value < 0.01) and two CAZymes, GH18 (a chitinase) and GH20 (hexosaminidase), were also more common in the host-associated genomes (panGWAS odds-ratio = 3.92; p-value < 0.001; **Supplementary Table 4**)). Taken together, our analysis of functional pathways suggests that the bacterial genus *Pseudoalteromonas* may be evolutionarily primed for symbiosis, as most lineages within this genus inherently possess genomic machinery that would provide nutritional benefits for animal hosts (e.g., the nearly universal ability to produce Vitamins B1, B2, B7, and the majority of the genus maintaining an additional pathway for Vitamin B12). Despite only being found in 50% of *Gammaproteobacteria*, Type II Secretion, which allows for nutrient absorption from the environment and promotes symbiosis in non-pathogenic bacteria (Cianciotto & White, 2017; Jani & Cotter, 2010; Tseng et al., 2009), was universal across the pangenome.

### Localization of Pseudoalteromonas near mucus glands in nematodes

We further investigated the physical location of host-associated *Pseudoalteromonas* bacteria using fluorescence *in-situ* hybridization (FISH) to visualize symbionts within the body of oncholaimid marine nematodes. Using nematode specimens collected from the Florida Keys, we used specific probes to detect *Pseudoalteromonas* (Eilers et al., 2000). Our FISH results are consistent with previous SEM observations of “rod-shaped” bacteria in the mouth of oncholaimid nematodes isolated from deep-sea hydrothermal vents (Bellec et al., 2018), which is the expected phenotype of *Pseudoalteromonas* bacterial cells. We observed *Pseudoalteromonas* bacteria only in the nematode mouth, forming biofilms on the teeth and walls of the buccal cavity (**Fig. 7**), and as biofilms lining the sides of the esophagus down to the esophageal bulb (**Fig. 7**), which was consistent with biofilm functional pathway prediction from genome analysis (described above). Past the esophageal bulb, host cells were stained blue by DAPI, but there were no positive results for *Pseudoalteromonas*. Importantly, *Pseudoalteromonas* bacteria were not present in the nematode gut or intestinal tract (**Fig. 7**), suggesting that marine nematodes do not digest these symbionts as a food source.

**Fig. 7:**
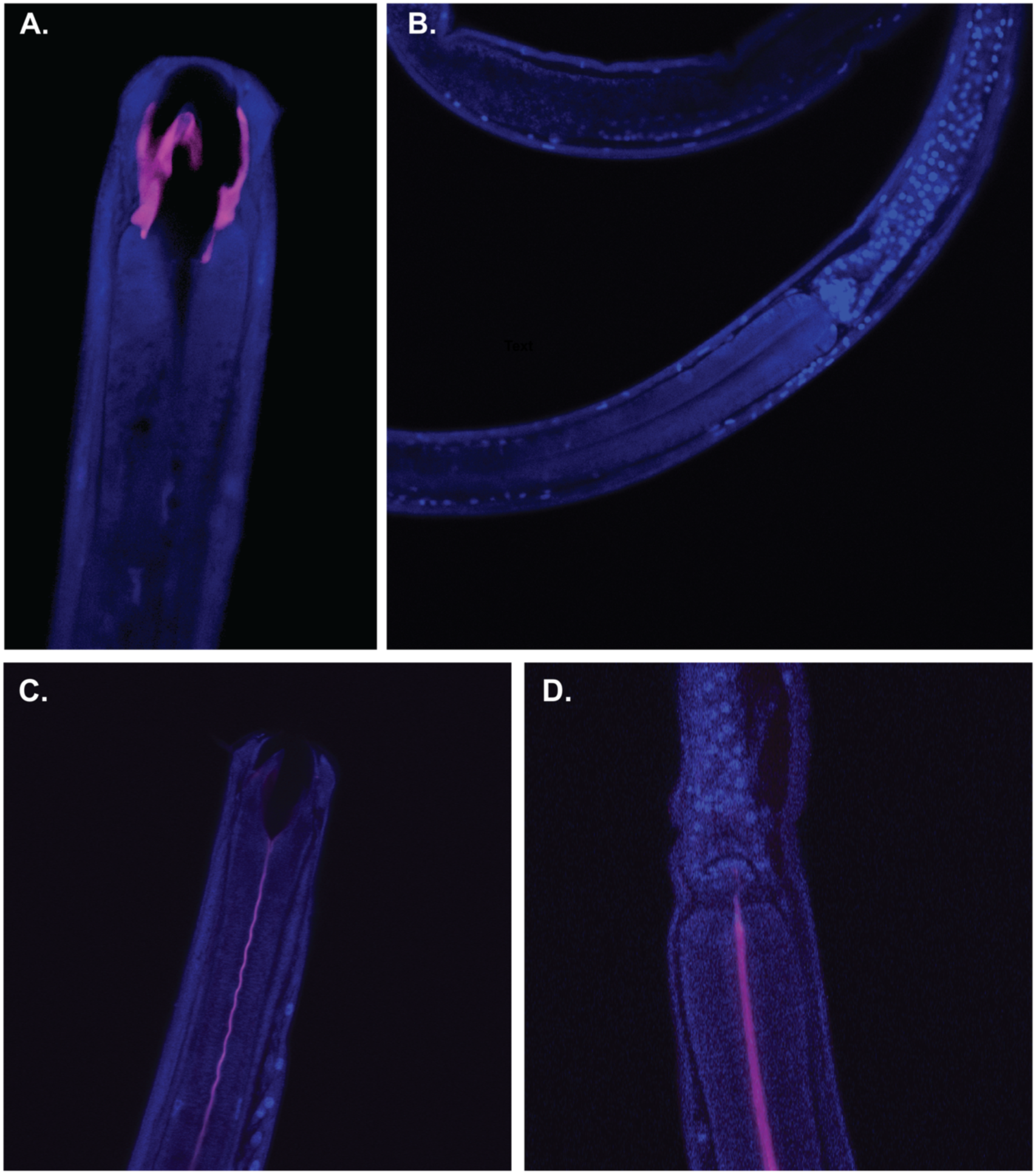
*Pseudoalteromonas* symbionts inhabit a physical niche in the mouth and esophagus of oncholaimid nematodes, localized near mucus excretions from the pharyngeal saliva glands. The blue indicates DAPI-stained nuclei of the nematode host (Nematoda, family Oncholaimidae). Pink color indicates the fluorescence of the *Pseudoalteromonas*-specific probes. There is no fluorescence of the *Pseudoalteromonas*-specific probes past the esophageal bulb of A and B) Oncholaimellus and B and C) *Metoncholaimus* nematodes, suggesting that host nematodes do not digest Pseudoalteromonas bacteria as a food source.

## Discussion

Our data broadly supports the existence of phylosymbiosis in marine invertebrate microbiomes, as evidenced through pangenomic analyses, host/bacterial phylogenetic topologies, the geographic consistency of symbiosis patterns (e.g., inclusive of Antarctic invertebrates), and FISH imaging which confirms the formation of host-associated biofilms and identifies a physical niche for novel *Pseudoalteromonas* symbionts. Our pangenomic study challenges the prior claim by Boscaro et al. (2022) that marine invertebrates lack phylosymbiosis, emphasizing the inadequacy of low-resolution 16S rRNA marker gene surveys for robustly assessing marine phylosymbiosis, and demonstrating the need for focused studies and phylogenomic analyses to evaluate evolutionary patterns. In our study of *Pseudoalteromonas*, phylogenetic comparisons of host/symbiont tree topologies exhibited patterns consistent with predictions for complex symbioses involving multiple hosts and bacterial lineages (Dismukes et al., 2022). By using genome-scale data from a single bacterial genus, we found substantial evidence of cophylogenetic signals between *Pseudoalteromonas* and marine invertebrates at the kingdom (Animalia), phylum (Nematoda, Mollusca, and Cnidaria), and family levels (Bivalvia and Faviidae) but weak or no evidence of phylogenetic congruence. Phylogenetic congruence is the most stringent method to assess cophylogeny, but such strict patterns are more likely to emerge in metazoan systems with vertically-acquired symbionts. In marine environments, horizontal transmission is the predominant method of symbiont acquisition (Breusing et al., 2022; Bright & Bulgheresi, 2010; Hauer et al., 2023), which often results in phylogenetic incongruence despite other scenarios that can lead to cophylogenetic signals, such as trait-matching and vicariance events (Dismukes et al., 2022; Perez-Lamarque & Morlon, 2024). Trait-based cophylogenetic signals, when the host interacts with bacterial groups that have a specific genomic trait, often cause phylogenetic incongruence, particularly in cases of bacterial groups that undergo substantial host-switching (Dismukes et al., 2022; Russo et al., 2018). Our phylogenomic and pangenomic evidence indicates that *Pseudoalteromonas* has an open pangenome which is associated with bacteria with a facultative mutualistic lifestyle and large effective population size (Dewar et al., 2024), which can further obscure cophylogenetic signals. In addition, at least five *Pseudoalteromonas* lineages appear to co-exist in marine nematode microbiomes, consistent with the emerging view that intra-host symbiont diversity is more prevalent than previously acknowledged (Ansorge et al., 2019; Gould et al., 2023). Our confirmation of phylosymbiosis between marine invertebrates and *Pseudoalteromonas*, a common but overlooked “core microbiome” taxon, suggests that host-bacterial codiversification in marine ecosystems is more common than currently understood, and that other “core microbiome” taxa contributing to phylosymbiosis signals in diverse marine invertebrates (Leasi et al., 2024; O’Brien et al., 2019, 2020; Tsang et al., 2024) should be the first target for deeper analyses that leverage whole bacterial genomes.

The host-associated lifestyle of *Pseudoalteromonas* is indicative of a nutritional mutualism symbiosis, where bacteria colonize specific physical niches on their animal hosts (e.g., forming biofilms in the nematode mouth and esophagus), outcompete other taxa, and stave off pathogens via production of antimicrobial compounds. In turn, *Pseudoalteromonas* lineages appear to provide their marine invertebrate hosts with food in the form of small organic molecules (simple carbohydrates and protein) as well as essential Vitamin B compounds, which animal hosts must obtain exogenously. Novel *Pseudoalteromonas* symbionts exhibit strong parallels with a number of other host/symbiont systems in terrestrial and marine ecosystems.

First, the features of host-associated *Pseudoalteromonas* appear to mirror those of another emerging *Gammaproteobacteria* symbionts in marine invertebrates, *Endozoicomonas*. Both taxa demonstrate extracellular aggregations of symbiont bacteria, enrichment of pathways for cycling and transport of carbohydrates and protein, enrichment of Vitamin B biosynthesis, and larger genomes with a high degree of genomic plasticity (including a high number of mobile genetic elements) which likely facilitates host switching and lifestyle flexibility in these bacterial genera (Bosi et al., 2017; Neave et al., 2016; Neave, Michell, et al., 2017; Pogoreutz et al., 2022). Pangenomes similar to *Pseudoalteromonas* and *Endozoicomonas* – containing a larger repertoire of accessory and unique genes – are associated with higher rates of horizontal gene transfer, which allows bacteria to rapidly gain and lose genes and increase their genetic diversity compared to closely related strains (S. L. Lim et al., 2025; McInerney et al., 2017), providing a competitive advantage for facultative symbiosis. Multiple *Endozoicomonas* are often recovered in association with a single coral host, further suggesting that niche partitioning amongst co-occurring symbiont strains may be typical for these extracellular, horizontally-acquired marine invertebrate symbionts (Neave et al., 2016; Neave, Michell, et al., 2017).

Second, the enrichment of Vitamin B biosynthesis and transport pathways in *Pseudoalteromonas* is analogous to terrestrial arthropod symbioses, where bacterial symbionts provide hosts with up to eight B Vitamin molecules that supplement the nutritionally-poor diet of blood- and sap-feeding host species (Buysse et al., 2021; Douglas, 2017). Bacterial supplies of B Vitamins are known to impact fitness, development, and reproduction in a range of invertebrate taxa (Croft et al., 2005; Douglas, 2017; Feng et al., 2024; Haçariz et al., 2021; Hickin et al., 2022). For nematodes in particular, bacterial B Vitamin production has been shown to enhance predatory behavior known as “surplus killing,” where nematode hosts kill an excess of prey that they do not consume (Akduman et al., 2020). Thus, genomic pathways enriched in host-associated *Pseudoalteromonas* may directly influence host growth, behavior, and reproduction, which in turn provides a direct benefit to extracellular symbionts by increasing the local availability of food and host-associated niches.

We hypothesize that *Pseudoalteromonas* bacteria may be reliant on the mucus of their invertebrate hosts, using mucus as a source of nutrition and/or protective agent that facilitates symbiosis. Invertebrate mucus has ancient evolutionary origins, and mucin-like glycoproteins or domains have been identified in fungi, parasites, and even viruses (Buscaglia et al., 2006; Freire et al., 2003; Lang et al., 2007; Wang et al., 2003), suggesting the possibility of host-symbiont co-evolution over long timescales. Animals produce mucus as a physical and chemical barrier that serves as the first line of defense against pathogen infection and disturbance (Glasl et al., 2016; Sheng & Hasnain, 2022; Shnit-Orland & Kushmaro, 2009), and many marine invertebrates utilize mucus to facilitate locomotion or adhesion, gather prey and detritus (Riemann & Helmke, 2002; Riemann & Schrage, 1978; Wilden et al., 2019), or enhance reproduction (e.g., as an antimicrobial-infused surface for laying eggs (Stabili, 2019)).

Furthermore, mucus properties are species-specific in marine invertebrates, exhibiting distinct nutritional profiles (e.g., varying ratios of carbohydrates, proteins, and lipids (Stabili, 2019; Stabili et al., 2019)) and often producing a high number of bioactive and antimicrobial compounds (with the mucus of polychaetes, corals, and sponges often targeted for natural products and drug discovery research (Li et al., 2023; Shnit-Orland & Kushmaro, 2009; Stabili, 2019; Stabili et al., 2019; Varijakzhan et al., 2021)). In corals, heterotrophic bacteria are known to quickly colonize mucus layers, and are capable of growing on mucus as their sole source of carbon (Pascal, 1982; Stabili, 2019), and other commensal bacteria such as *Akkermansia*, *Bacteroides*, and *Vibrio* are known to feed on or degrade host mucus using mucolytic enzymes (Crowther et al., 1987; Sheng & Hasnain, 2022). Experimental work has also shown that the mucus of oncholaimid nematodes promotes the exclusive growth of *Pseudoalteromonas* bacteria, and oncholaimid nematodes are known to excrete copious amounts of mucus from the pharyngeal salivary glands where our FISH images indicated bacterial biofilms were localized (Moens et al., 2005; Riemann & Schrage, 1978). Notably, oncholaimid nematodes are also known to rapidly pump seawater through their intestinal tract (Moens et al., 1999), and can directly uptake dissolved organic carbon compounds such as glucose and acetate through their intestinal wall (Chia & Warwick, 1969; Riemann et al., 1990), suggesting that nematode hosts may rapidly ingest and assimilate bacterial B vitamins and small organic molecules produced by *Pseudoalteromonas* symbiont biofilms.

Physical attachment to the mucus layer of invertebrate hosts is also thought to maximize the activity of bioactive substances and provide a competitive advantage for bacteria (e.g., preventing dilution of antimicrobial compounds into the surrounding water (O’Brien et al., 2019; Shnit-Orland & Kushmaro, 2009)). The localization of *Pseudoalteromonas* in the nematode mouth and esophagus may also point towards multiple successional stages of bacterial colonization, or species-specific symbioses where the physical host niche varies across different oncholaimid nematodes; however, further experimental work would be needed to confirm these hypotheses. We hypothesize that invertebrate mucus is the specific trait driving the cophylogenetic signals reported in this study, with symbiosis and host-switching determined by the molecular composition and antimicrobial properties of invertebrate host species’ mucus.

Over evolutionary timescales, invertebrate mucus may have served as a protected microhabitat and consistent source of food, encouraging the establishment of symbiosis and codiversification of invertebrate-bacterial partnerships in marine environments worldwide.

## Materials and Methods

Unless otherwise specified, all bioinformatics tools and computational algorithms were run using default parameters. All scripts utilized during this study have been made available on GitHub: https://github.com/BikLab/Pseudoalteromonas.

### Isolation and sequencing of nematode-associated Pseudoalteromonas

To isolate and culture nematode-associated bacteria, marine nematodes from the family Oncholaimidae were isolated from sediment samples collected from Tybee Island, GA, UGA in 2020 (De Santiago et al., 2025). Following the methods of Schuelke et al. (2018), marine nematodes were isolated using the decantation-flotation method (Danovaro, 2009), and sediment was washed over 45 μm sieves using artificial seawater (Instant Ocean® Spectrum Brands, Inc., USA) prepared with Milli-Q ultrapure water. Nematodes were individually picked under a dissection microscope (Olympus SZX16, Olympus Corporation, Tokyo, Japan) and transferred to sterile artificial seawater on a temporary slide mount. Nematodes were then taxonomically identified under an Olympus BX63 compound microscope to the genus level following the World Register of Marine Species (WoRMS) database (WoRMS Editorial Board, 2025) using specialized morphological keys for marine nematodes (Platt et al., 1983, 1985; Platt & Warwick, 1988; Warwick et al., 1998). Nematodes were quickly rinsed in molecular-grade water, placed in a microcentrifuge tube with sterile artificial seawater (Instant Ocean® Spectrum Brands, Inc., USA), and immediately shipped on ice to the University of Southern California, where the sample underwent dilution-to-extinction isolation and were later genome-sequenced, as described (De Santiago et al., 2025).

### Isolation of marine nematodes for single-worm metagenomics

To identify whether *Pseudoalteromonas* constitutes part of the core nematode microbiome, marine nematodes were isolated from diverse habitats and locations to undergo short-read metagenomic sequencing. Nematodes were isolated from frozen samples previously collected as part of other studies (representing geographic locations in Bodega Bay, CA, Tybee Island, GA, and the continental shelf around Antarctica). As described above, nematodes were isolated from sediments using a decantation-flotation method (Danovaro, 2009). Samples were washed over a 45 μm sieve using artificial seawater (Instant Ocean® Spectrum Brands, Inc., USA) prepared with Milli-Q ultrapure water. Nematodes were individually picked under a dissection microscope and transferred to sterile artificial seawater on a temporary slide mount. Individual worms were then taxonomically identified to the family or genus level following the World Registers of Marine Species (WoRMS) database (WoRMS Editorial Board, 2025) using specialized morphological keys for marine nematodes (Platt et al., 1983, 1985; Platt & Warwick, 1988; Warwick et al., 1998) as described above. Worms were quickly rinsed in molecular-grade water to remove transient environmental microbes attached to the cuticle before being stored in microcentrifuge tubes with either 20 μL of water or 10 μL REPLI-g Single Cell Cryo-protect Reagent (QIAGEN). One nematode specimen was placed in each tube to characterize the microbiome associated with each worm. Lab workspaces and equipment were sterilized with 70% ethanol before and between samples. Nematodes from Antarctica were frozen in DESS (Yoder et al., 2006) and transferred to 10 μL REPLI-g Single Cell Cryo-protect Reagent (QIAGEN) prior to DNA extraction and amplification.

Worms initially placed in 20 μL of molecular-grade water underwent DNA extraction using the E.Z.N.A® MicroElute Genomic DNA Kit following manufacturer’s protocol. Worms in 10 μL REPLI-g Single Cell Cryo-protect Reagent (QIAGEN) underwent DNA lysis and multiple displacement amplification (MDA) using the REPLI-g Advanced DNA Single Cell Kit (QIAGEN) with a modified protocol to accommodate storage of the worms (which were stored in 10 μL of REPLI-g Single Cell Cryo-protect Reagent compared to the manufacturer’s recommended 4 μL) and to target the lysis of bacterial cells. A total of 6 μL of Buffer D2, containing 0.5 μL of 1 M DDT and 5.5 μL of reconstituted buffer DLB, was added to the nematodes stored in 10 μL REPLI-g Single Cell Cryo-Protect Reagent (QIAGEN) to lyse the nematode-associated bacterial cells. Tubes were vortexed and centrifuged to briefly mix the samples. The reaction mixture was incubated in a Thermomixer heated shaker block (Eppendorf, Hamburg, Germany) at 65°C for 10 min. A total of 6 μL of Stop Solution was added to each tube. Amplification reactions were 50 μL total reaction, containing 9 μL of water, 29 μL of REPLI-g advanced sc Reaction Buffer, 2μL of REPLI-g DNA polymerase, and 10 μL of denatured DNA. The samples were incubated in a T1000 Thermocycler (Bio-Rad) at 30°C for 2 hours with a lid temperature of 70°C, followed by 65°C for 2 min to inactivate the amplification reaction. Samples that underwent successful amplification or had high DNA biomass during the initial lysis step were sent for Illumina library preparation and Illumina sequencing using the NextSeq Platform (300 bp PE) at either the Department of Energy Joint Genome Institute or the Georgia Genomics and Bioinformatics Core at the University of Georgia. A subset of nematode genera, representing four distinct families, was included in this study (**Supplementary Table 2**).

### Constructing the Pseudoalteromonas pangenome

A total of 236 *Pseudoalteromonas* genomes, including all 234 high-quality genomes available in RefSeq (completeness >95%; contamination <5%) as of August 2024 and the two novel nematode-associated genomes generated as part of this study, were used to construct the *Pseudoalteromonas* pangenome (**Supplementary Table 1**). The collection consisted of 91 genomes from bacteria isolated from seawater (“free-living” isolates) and 145 genomes isolated from marine invertebrates (“host-associated” isolates). We performed gene-calling and annotation using PROKKA v1.14.5 (Seemann, 2014) with UniProt (Acids research & 2023, 2023) as a protein reference database. PIRATE v1.0.5 (Bayliss et al., 2019) was used to identify orthologous genes and construct the pangenome using the default percent identity thresholds recommended by the developers. A custom RScript (available on GitHub) was used to group orthologous gene families into core (genes present in >95% of the genomes), accessory (genes present in <95% and found in more than one genome), and unique genes (genes present in only one genome). A Pearson’s Chi-squared statistical test was used to test if there was a relationship between the frequency of genes (core, accessory, and unique) and lifestyle. To determine whether *Pseudoalteromonas* exhibited an open or closed pangenome, we calculated gene accumulation curves of the gene families using a custom RScript (also available on GitHub at the repository linked above). The pangenome was designated as ‘open’ if new gene families were identified as new genomes were added (i.e., the accumulation curve did not reach an asymptote). The pangenome was classified as “closed” if no new gene families were found (i.e., the accumulation curve reached an asymptote).

### Constructing the Pseudoalteromonas phylogenomic tree and population structure

To construct a pangenome-based phylogeny of *Pseudoalteromonas*, PIRATE v1.0.5 (Bayliss et al., 2019) was used to identify and individually align the 1,689 core genes using MAFFT v7.5.20 (Katoh & Standley, 2013) and subsequently concatenate them into a nucleotide core alignment. The core alignment was used to construct a maximum-likelihood phylogenetic tree with RAxML v8.2.12 (Stamatakis, 2014) using the General Time Reversible (GTR) model with GAMMA-distributed rates across sites and 10,000 bootstraps. The dRep v3.4.2 (Olm et al., 2017), with the fastANI algorithm (Jain et al., 2018), was used to compare genomic relatedness and cluster genomes with 95% similarity into phylogenetic groups.

To identify whether the accessory genes were shared between phylogroups or lifestyles, a genome-genome network was constructed. The PIRATE presence/absence matrix was processed using the GraPPLE toolkit (Harling-Lee et al., 2022) to explore the accessory pangenome. The presence/absence matrix was filtered to include only the accessory genes present in more than 5% of the bacterial genomes. Core and unique gene families were excluded from this analysis. The presence/absence matrix was converted into a binary format using the GraPPLE script ‘gene_matrix_to_binary.py’. The GraPPLE script ‘pw_similarity.py’ was run to calculate pairwise similarities between genomes to create a genome-genome network and compare genomic similarities between lifestyle (host-associated vs. free-living) and phylogroups. Finally, the genome-genome network graphs were visualized using the Graphia v4.2 platform (Freeman et al., 2022).

### Evaluating phylosymbiosis among Pseudoalteromonas spp. and marine invertebrate hosts

Following methods used in previous studies of bacterial-animal phylosymbiosis (Gregor et al., 2022), midrooted phylogenetic trees were constructed using the core alignments for Mollusca, Cnidaria, and Porifera-associated *Pseudoalteromonas*. This analysis was restricted to these phyla due to insufficient genomic representatives across other invertebrate phyla. The alignments were used to construct a maximum-likelihood phylogenetic tree with RAxML v8.2.12 (Stamatakis, 2014) using the General Time Reversible (GTR) model with GAMMA-distributed rates across sites and 10,000 bootstraps. The host phylogenies (i.e., Mollusca, Cnidaria, and Porifera) were inferred using the TimeTree Database v5 (Kumar et al., 2022).

Cophylogenetic signal was defined as a non-random association between the host and symbiont phylogenies (Dismukes et al., 2022; Perez-Lamarque & Morlon, 2024). Phylogenetic congruence is a specific example of a cophylogenetic signal characterized by shared patterns of codiversification between the host and their symbiont (Dismukes et al., 2022; Perez-Lamarque & Morlon, 2024). Cophylogenetic signals were tested using two methods: 1) PACo (i.e., a Procrustean Approach to test for Congruence of two phylogenetic trees) using the paco R package and 2) using the Generalized Robinson–Foulds (RF) metric (Böcker et al., 2013) implemented in the TreeDist R package. Cophylogenetic signal was determined if the PACo analysis, with 1000 permutations, was significant (p-value < 0.05). Evidence of cophylogenetic signal was determined using the Generalized RF metric and PACo Coefficient of Determination (R^2^). R^2^ was calculated from the Procrustes squared sums (R^2^ =1 - m^2^). A high R^2^ (1) indicates perfect phylogenetic congruence. Following the recommendations of Perez-Lamarque & Morlon (2024), phylogenetic congruence was determined if the PACo analysis was significant with an R^2^ >0.25. Phylogenetic congruence was further validated using the generalized Robinson-Foulds (RF) metric.

To further visualize phylogenetic and geographic patterns seen in nematode-associated *Pseudoalteromonas* strains, we used a read-mapping approach whereby single-worm metagenomes (e.g., low complexity metagenomes containing only a single host genome and its associated microbiome community) were mapped to the *Pseudoalteromonas* pangenome. The quality of the sequences was visualized using FastQC (Andrews & Others, 2010) and summarized using MultiQC (Ewels et al., 2016). Sequences were quality trimmed and underwent adaptor removal using Trimmomatic (Bolger et al., 2014) with a sliding window (4:15) and a minimum length of 36 bp. A k-mer approach, implemented using Kraken2 (Wood et al., 2019), was used to identify and extract bacterial reads from the nematode/microbiome metagenomic dataset. The quality-controlled sequences were mapped to *Pseudoalteromonas* genomes using the Read Recruitment Analysis Pipeline (RRAP) (Kojima et al., 2022). RRAP normalizes the number of mapped reads by sequencing depth and genome size by using the RPKM (reads per kilobase [of the genome] per million [bases of recruited sequences]) method. An RPKM threshold of 0.1 was implemented to account for false positives. The results were visualized as a heatmap and phylogenetic framework using the ggtree (Yu et al., 2017) package in R. The PACo analysis was used to test for cophylogenetic signals between the nematode phylogeny and the hierarchical clustering of the read mapping dissimilarity matrix using the Bray-Curtis distance matrix.

### Identifying metabolic pathways and secondary metabolites across the pangenome

METABOLIC-G v4.0 (Zhou et al., 2022) was run with the full KOfam database (Aramaki et al., 2020) to predict the metabolic and biogeochemical traits of each bacterial genome included in this study. A multiSMASH (https://github.com/zreitz/multismash) Snakemake pipeline was used to summarize the secondary metabolite biosynthesis gene clusters predicted by the antiSMASH software (Blin et al., 2023). The presence/absence of the METABOLIC and antiSMASH results were imported into R for downstream data analysis. METABOLIC and antiSMASH results were visualized in a phylogenetic framework using the ggtree package (Yu et al., 2017) in R. The Scoary2 software (Roder et al., 2024) was used to conduct a pangenomic genome-wide association study (GWAS) to identify genes (using the pangenomic presence/absence table from PIRATE) and functions (using the METABOLIC and antiSMASH results) that co-occur with the bacterial lifestyles (i.e., host-associated or free-living) and phylogroups. Scoary2 determines significant trait-associated genes using a Fisher’s test (p-value < 0.05) with a Bonferroni adjustment using a family-wise error rate (FWER) cutoff of 0.99 to reduce the false positive discovery rate. Hypothetical proteins found to be more common in host-associated genomes were annotated by Gaia, a novel Artificial Intelligence tool developed by Tatta Bio, that integrates several genomic and structural features to predict functions of novel proteins (https://gaia.tatta.bio/) (Jha et al., 2025).

### Visualizing *Pseudoalteromonas* bacteria in nematodes using FISH

Oncholaimidae nematodes collected from the Florida Keys were processed using a modified protocol previously used for fluorescent *in situ* hybridization (FISH) of nematode-associated bacteria (Bellec et al., 2019). Nematodes belonging to the family Oncholaimidae were fixed for 2h in 3% formaldehyde with sterile artificial seawater (Instant Ocean® Spectrum Brands, Inc., USA), followed by three rinses in a 1:1 solution of PBS:Sterile seawater. The samples were stored in 1:1 storage buffer (100% ethanol:2X PBS) and kept frozen at -20°C until they were used. Nematodes were rinsed in 30% formamide buffer and hybridized in 150µL hybridization buffer [0.9M NaCl, 0.02M Tris-HCl (pH 8.0), 0.01% SDS, 2mL blocking reagent [5% Bovine Serum Albumin (BSA in 0.5% PBS-Tween 0.5%) and 95% 7.4 pH PBS], and 30% formamide] at 46°C for 3hr with 15µL (8µM) of each probe: universal bacterial probe (5’-GCTGCCTCCCGTAGGAGT-3’; CY-3; (Amann et al., 1990)), *Pseudoalteromonas*-specific probe (5’-CCCCTTTGGTCCGTAGAC-3’; CY-5; (Eilers et al., 2000)), and nonsense probe (5’-ACTCCTACGGGAGGCAGC-3’; FAM; (Wallner et al., 1993)). The nematodes were rinsed three times in washing buffer [0.046M NaCl, 0.02M Tris-HCl (pH 80), 0.005 M EDTA, 0.01% SDS] for 30 min at 46°C for a total of 1.5 hrs. The hybridized specimens were mounted on a slide in SlowFade^TM^ Gold Antifade reagent (Invitrogen) containing DAPI and observed using the Zeiss LSM 880 Confocal Microscope system. The Zen 2.3 imaging software was used for image acquisition and processing.

## ACKNOWLEDGEMENTS

Funding for this study was provided by the Gordon and Betty Moore Foundation (Symbiosis in Aquatic Systems Initiative, grant #9326), and a National Science Foundation CAREER award (DEB-2144304) and Antarctic Program award (OPP-2132641) to HMB at UGA. Research support for ADS was provided by the University of Georgia Research Foundation and the National Institute of General Medical Sciences of the National Institute of Health under award number 1T32GM142623. We acknowledge the Joint Genome Institute (JGI, Proposal ID: 505025) and the Georgia Genomics and Bioinformatics Core (GGBC, UGA, RRID:SCR_010994) for metagenomic sequencing of single-worm isolates and the Georgia Advanced Computing Resource Center (GACRC) at UGA (https://gacrc.uga.edu/) for computational resources that have contributed to the results in this publication. The work (proposal: 10.46936/10.25585/60001240) conducted by the U.S. Department of Energy Joint Genome Institute (https://ror.org/04xm1d337), a DOE Office of Science User Facility, is supported by the Office of Science of the U.S. Department of Energy operated under Contract No. DE-AC02-05CH11231. We also acknowledge the Biomedical Microscopy Core (BMC) at UGA for their assistance and support in generating the FISH micrographs for the study. We thank Kevin Kocot for providing samples from the Antarctic cruise NBP-20-10.

## Data Availability

All the genomes used in this study are publicly available in the NCBI RefSeq (see **Supplementary Table 1** for all associated accession codes). Raw metagenomic sequences generated by the Georgia Genomics and Bioinformatics Core (GGBC) and the Joint Genomic Institute (JGI, Proposal ID: 505025) were deposited in the NCBI Sequence Read Archive (BioProject: XXXXXX).

## Code Availability

All scripts and files required for the reproducibility of all analyses conducted in this study are available via GitHub (https://github.com/BikLab/Pseudoalteromonas)

## Supplemental Material

**Extended Data Fig. 1: Average nucleotide identity reveals a higher diversity within the genus Pseudoalteromonas than previously estimated.** A higher fastANI score correlates with high genetic similarity between clades. A minimum of 95% fastANI was used to place bacterial isolates in the same phylogroup.

**Extended Data Fig. 2. Additional evidence for cophylogenetic signals between *Pseudoalteromonas* and marine invertebrates.** Phylogenetic trees of *Pseudoalteromonas* (top) and the hosts (bottom) were used to test for cophylogeny across the Kingdom (Animalia). Topological trees (top) of *Pseudoalteromonas* were constructed with RAxML (Stamatakis, 2014) using the General Time Reversible (GTR) model with GAMMA-distributed rates across sites and 100 bootstraps. Host cladograms (bottom) were generated via TimeTree (Kumar et al., 2022). Cophylogeny signal was tested using PACo, and phylogenetic congruence was tested using the Generalized Robinson-Foulds metric (**Table 1**).

**Extended Data Fig. 3: Host-associated genomes contain distinct sets of genes that enable host-bacterial interactions.** A) Midrooted phylogenetic tree of the *Pseudoalteromonas* pangenome. The tips indicate whether the isolate was a free-living (blue circle) or host-associated (red circle) strain, and the branch color indicates the phylogroup. B) Summary of the accessory genes that were more common in the host-associated pigmented (orange) or host-associated nonpigmented (purple) bacterial genomes. The results of the SCOARY2 panGWAS and a summary of the gene products can be found in **Supplementary Table 3**.

**Supplementary Table 1: Summary of the genomes included in this study.** The lifestyle of the isolated strain (host-associated vs. free-living), the phyla of the host it was isolated from, the completeness and contamination (based on CheckM), and the AccessionID. The two genomes isolated from marine nematodes in this study are highlighted in yellow.

Supplementary Table 2: Summary of single-worm metagenomic samples used to identify the prevalence of *Pseudoalteromonas* in the microbiome of marine nematodes. Metadata includes the habitat that the worms were isolated from, the genus-level identification of the worm, the genome amplification protocol, and the date they were collected from.

**Supplementary Table 3: Accessory genes more commonly found in host-associated genomes.** A summary of the genes more commonly found in host-associated (t+) pigmented (orange) and nonpigmented (purple) genomes. The gene family ID is in bold, followed by a brief description of the product. The presence and absence of a function or gene is indicated by (g+) or (g-), respectively. Functional predictions of hypothetical proteins by Gaia, an AI agent developed by Tatta Bio that integrates several genomic and structural features to predict functions of novel proteins, are in red. An odds-ratio greater than one indicates that the gene/function is more commonly associated with the host-associated lifestyle strains.

**Supplementary Table 4: Metabolic functions and biogeochemical traits more commonly found in host-associated genomes.** A summary of the metabolic functions and enzymes associated with host-associated (t+) pigmented (orange) and nonpigmented (purple) genomes. The name of the enzyme or function is in bold, followed by a brief description of its function. The presence and absence of a function or gene is indicated by (g+) or (g-), respectively. An odds-ratio greater than one indicates that the gene/function is more commonly associated with the host-associated lifestyle strains.

