## Supplementary Table 1 for "*Pseudoalteromonas* is a novel symbiont of marine invertebrates that exhibits broad patterns of phylosymbiosis"

**Supplementary Table 1: Summary of the genomes included in this study.** The lifestyle of the isolated strain (host-associated vs. free-living), the phyla of the host it was isolated from, the completeness and contamination (based on CheckM), and the AccessionID. The two genomes isolated from marine nematodes in this study are highlighted in yellow.

| **SampleID** | **AccessionID** | **Phylogroup** | **Completeness** | **Contamination** | **Lifestyle** | **Host Phylum** |
| --- | --- | --- | --- | --- | --- | --- |
| **P_1131a** | GCF_001401875.1 | PG_44 | 100 | 0.57 | Host-associated | Cnidaria |
| **P_1161b** | GCF_001401785.1 | PG_35 | 99.95 | 0.25 | Host-associated | Cnidaria |
| **P_125** | GCF_001401805.1 | PG_44 | 99.75 | 0.82 | Host-associated | Cnidaria |
| **P_126** | GCF_001401795.1 | PG_30 | 100 | 0.73 | Host-associated | Cnidaria |
| **P_130** | GCF_001400005.1 | PG_45 | 100 | 0.38 | Host-associated | Cnidaria |
| **P_17a** | GCF_001401855.1 | PG_44 | 100 | 0.51 | Host-associated | Cnidaria |
| **P_1CM17D** | GCF_023161805.1 | PG_45 | 100 | 0.25 | Host-associated | Cnidaria |
| **P_ACER1** | GCF_024124445.1 | PG_29 | 99.61 | 1.22 | Host-associated | Cnidaria |
| **P_APAL1** | GCF_021532475.1 | PG_26 | 99.70 | 1.07 | Host-associated | Cnidaria |
| **P_2CM32C** | GCF_023161925.1 | PG_43 | 100 | 0.13 | Host-associated | Porifera |
| **P_2CM36K** | GCF_023161865.1 | PG_34 | 99.95 | 0.27 | Host-associated | Chordata |
| **P_B95** | GCF_030125375.1 | PG_03 | 99.95 | 2.63 | Host-associated | Cnidaria |
| **P_BZA4** | GCF_040104395.1 | PG_30 | 99.70 | 0.63 | Host-associated | Cnidaria |
| **P_P94** | GCF_030135505.1 | PG_02 | 99.70 | 0.41 | Host-associated | Cnidaria |
| **P_piratica_OCN003** | GCF_000788395.1 | PG_55 | 100 | 1.31 | Host-associated | Cnidaria |
| **P_A22** | GCF_018141345.1 | PG_15 | 99.12 | 0.23 | Host-associated | Arthropoda |
| **P_A25** | GCF_009176705.1 | PG_22 | 99.75 | 0.23 | Free-living | Seawater |
| **P_AC163** | GCF_000497935.1 | PG_40 | 95.75 | 1.3 | Host-associated | Porifera |
| **P_R96** | GCF_030125445.1 | PG_06 | 99.96 | 0.76 | Host-associated | Cnidaria |
| **P_agarivorans_DSM14585** | GCF_002310855.1 | PG_45 | 99.20 | 0.25 | Free-living | Seawater |
| **P_agarivorans_NW4327** | GCF_000508785.1 | PG_45 | 100 | 3.79 | Host-associated | Porifera |
| **P_agarivorans_S816** | GCF_000363985.1 | PG_45 | 100 | 0.51 | Free-living | Seawater |
| **P_aliena_EH1** | GCF_001999225.1 | PG_38 | 100 | 0.62 | Free-living | Seawater |
| **P_aliena_SW19** | GCF_014905615.1 | PG_38 | 100 | 0.25 | Free-living | Seawater |
| **P_amylolytica_JW1** | GCF_001854605.1 | PG_21 | 99.49 | 0.54 | Free-living | Seawater |
| **P_2CM37A** | GCF_023161885.1 | PG_45 | 100 | 0.25 | Host-associated | Cnidaria |
| **P_APM04** | GCF_022809815.1 | PG_43 | 99.75 | 0.13 | Free-living | Seawater |
| **P_arctica_A3712** | GCF_000238395.3 | PG_42 | 99.75 | 0.38 | Free-living | Seawater |
| **P_arctica_NECBIFX0059** | GCF_012844465.1 | PG_41 | 100 | 0.13 | Host-associated | Mollusca |
| **P_arctica_NECBIFX2020001** | GCF_012933495.1 | PG_42 | 100 | 0.13 | Host-associated | Mollusca |
| **P_arctica_NECBIFX2020017** | GCF_012935355.1 | PG_42 | 100 | 0.13 | Host-associated | Echinodermata |
| **P_aurantia_208** | GCF_014858725.1 | PG_20 | 100 | 0.48 | Free-living | Seawater |
| **P_aurantia_S3788** | GCF_005886875.1 | PG_18 | 99.92 | 0.44 | Free-living | Seawater |
| **P_aurantia_S3790** | GCF_005887355.1 | PG_18 | 99.66 | 0.69 | Free-living | Seawater |
| **P_aurantia_S3895** | GCF_005887285.1 | PG_18 | 99.92 | 0.51 | Free-living | Seawater |
| **P_2CM41L** | GCF_023161905.1 | PG_45 | 100 | 0.25 | Host-associated | Cnidaria |
| **P_flavipulchra_SCSIO43202** | GCF_024746855.1 | PG_15 | 100 | 0.41 | Host-associated | Cnidaria |
| **P_SCSIO43088** | GCF_023650795.1 | PG_30 | 98.05 | 0.32 | Host-associated | Cnidaria |
| **P_SCSIO43101** | GCF_023650775.1 | PG_27 | 99.95 | 0.46 | Host-associated | Cnidaria |
| **P_SCSIO43201** | GCF_023716725.1 | PG_13 | 99.75 | 1.22 | Host-associated | Cnidaria |
| **P_BMB** | GCF_001709235.1 | PG_15 | 99.47 | 0.34 | Host-associated | Ctenophora |
| **P_Cn537** | GCF_021532495.1 | PG_29 | 99.95 | 0.76 | Host-associated | Cnidaria |
| **P_CNAT218** | GCF_021530715.1 | PG_16 | 99.37 | 0.97 | Host-associated | Cnidaria |
| **P_CNAT2181** | GCF_022347865.1 | PG_16 | 99.37 | 0.97 | Host-associated | Cnidaria |
| **P_C2R02** | GCF_018860845.1 | PG_57 | 98.70 | 3.70 | Free-living | Seawater |
| **P_citrea_DSM8771** | GCF_000238375.3 | PG_19 | 100 | 0.25 | Free-living | Seawater |
| **P_citrea_S2231** | GCF_005887475.1 | PG_17 | 99.92 | 0.65 | Host-associated | Arthropoda |
| **P_citrea_S2233** | GCF_005887445.1 | PG_17 | 99.92 | 0.79 | Host-associated | Arthropoda |
| **P_CNAT241** | GCF_040104515.1 | PG_16 | 99.37 | 0.97 | Host-associated | Cnidaria |
| **P_bablabjr011** | GCF_016464045.1 | PG_24 | 99.95 | 0.51 | Host-associated | Cnidaria |
| **P_DL2H1** | GCF_021532565.1 | PG_15 | 99.49 | 0.39 | Host-associated | Cnidaria |
| **P_DL2H22** | GCF_021532535.1 | PG_09 | 99.96 | 0.25 | Host-associated | Cnidaria |
| **P_DL2H6** | GCF_021530725.1 | PG_15 | 99.13 | 0.39 | Host-associated | Cnidaria |
| **P_shioyasakiensis_28** | GCF_024105465.1 | PG_30 | 97.50 | 0.34 | Host-associated | Cnidaria |
| **P_BZB3** | GCF_040104455.1 | PG_50 | 98.69 | 0.51 | Host-associated | Cnidaria |
| **P_Ps84H4** | GCF_024124645.1 | PG_26 | 99.70 | 1.25 | Host-associated | Cnidaria |
| **P_CO109Y** | GCF_004208935.1 | PG_30 | 99.95 | 0.53 | Host-associated | Cnidaria |
| **P_CO133X** | GCF_004208945.1 | PG_25 | 99.95 | 0.13 | Host-associated | Cnidaria |
| **P_CO302Y** | GCF_004209095.1 | PG_25 | 99.95 | 0.13 | Host-associated | Cnidaria |
| **P_CO325X** | GCF_004208955.1 | PG_16 | 99.49 | 0.72 | Host-associated | Cnidaria |
| **P_CO342X** | GCF_004208915.1 | PG_15 | 100 | 0.16 | Host-associated | Cnidaria |
| **P_CO348** | GCF_004208845.1 | PG_15 | 100 | 0.70 | Host-associated | Cnidaria |
| **P_D15MCD2** | GCF_038562745.1 | PG_30 | 99.62 | 0.77 | Free-living | Seawater |
| **P_distincta_16SW7** | GCF_005877035.1 | PG_40 | 100 | 0.38 | Free-living | Seawater |
| **P_distincta_22A13** | GCF_028414575.1 | PG_40 | 100 | 0.13 | Host-associated | Echinodermata |
| **P_distincta_ATCC700518** | GCF_000814675.1 | PG_40 | 99.75 | 0.13 | Host-associated | Porifera |
| **P_distincta_KMM3548** | GCF_014918315.1 | PG_40 | 98.70 | 0.13 | Host-associated | Cnidaria |
| **P_distincta_LMG14908** | GCF_030710225.1 | PG_40 | 100 | 0.13 | Host-associated | Mollusca |
| **P_B7P1** | GCF_040104415.1 | PG_30 | 99.70 | 0.98 | Host-associated | Cnidaria |
| **P_CNC920** | GCF_022347875.1 | PG_16 | 99.37 | 0.72 | Host-associated | Cnidaria |
| **P_MM172** | GCF_022347945.1 | PG_16 | 99.37 | 0.97 | Host-associated | Cnidaria |
| **P_donghaensis_HJ51** | GCF_003515105.1 | PG_24 | 99.95 | 1.01 | Free-living | Seawater |
| **P_DSM17587** | GCF_000238335.3 | PG_43 | 100 | 0.13 | Free-living | Seawater |
| **P_EB27** | GCF_001974875.1 | PG_41 | 99.75 | 0.13 | Host-associated | Porifera |
| **P_ECSMB14103** | GCF_000813065.1 | PG_43 | 100 | 0.13 | Host-associated | Mollusca |
| **P_EKP108** | GCF_014534795.1 | PG_40 | 100 | 0.38 | Free-living | Seawater |
| **P_espejiana_DSM9414** | GCF_002221525.1 | PG_46 | 99.45 | 0.51 | Free-living | Seawater |
| **P_FE4** | GCF_030297135.1 | PG_27 | 99.95 | 0.13 | Host-associated | Echinodermata |
| **P_flavipulchra_ATCCBAA314** | GCF_000814665.1 | PG_15 | 99.75 | 1.30 | Free-living | Seawater |
| **P_flavipulchra_LMG20361** | GCF_014596995.1 | PG_15 | 95.08 | 0.32 | Free-living | Seawater |
| **P_flavipulchra_NCIMB2033** | GCF_014858715.1 | PG_15 | 99.75 | 0.90 | Free-living | Seawater |
| **P_flavipulchra_S1925** | GCF_005886245.1 | PG_15 | 100 | 1.17 | Host-associated | Arthropoda |
| **P_BZK2** | GCF_014349195.1 | PG_29 | 99.99 | 0.61 | Host-associated | Cnidaria |
| **P_fuliginea_KMM216** | GCF_000690055.1 | PG_39 | 99.75 | 0.57 | Host-associated | Chordata |
| **P_fuliginea_S2292** | GCF_000967705.1 | PG_39 | 99.75 | 0.57 | Host-associated | Porifera |
| **P_G4** | GCF_028875915.1 | PG_54 | 98.57 | 1.54 | Host-associated | Arthropoda |
| **P_galatheae_S4498** | GCF_005886105.2 | PG_14 | 99.75 | 0.67 | Host-associated | Arthropoda |
| **P_gelatinilytica_NH153** | GCF_001641615.1 | PG_28 | 99.95 | 0.65 | Free-living | Seawater |
| **P_Hal040** | GCF_037162425.1 | PG_28 | 95.03 | 0.41 | Host-associated | Porifera |
| **P_HMSA03** | GCF_002289345.1 | PG_15 | 97.55 | 0.19 | Host-associated | Mollusca |
| **P_JC28** | GCF_013319255.1 | PG_15 | 99.49 | 0.38 | Host-associated | Mollusca |
| **P_JC3** | GCF_017168135.1 | PG_15 | 99.75 | 1.36 | Host-associated | Arthropoda |
| **P_JC31** | GCF_030291895.1 | PG_15 | 99.75 | 1.36 | Host-associated | Arthropoda |
| **P_JW3** | GCF_001854555.1 | PG_21 | 99.49 | 0.54 | Free-living | Seawater |
| **P_kknpp56** | GCF_018598885.1 | PG_34 | 99.77 | 0.76 | Free-living | Seawater |
| **P_KMM661** | GCF_002221505.1 | PG_37 | 99.89 | 0.54 | Host-associated | Mollusca |
| **P_lipolytica_CSB02KR** | GCF_001704875.1 | PG_24 | 99.95 | 0.48 | Host-associated | Echinodermata |
| **P_lipolytica_LMEB39** | GCF_014925285.1 | PG_24 | 99.87 | 0.23 | Free-living | Seawater |
| **P_luteoviolacea_2ta16** | GCF_000495575.1 | PG_01 | 98.94 | 1.70 | Host-associated | Cnidaria |
| **P_M1318928** | GCF_022398505.1 | PG_27 | 99.70 | 0.41 | Host-associated | Mollusca |
| **P_M1400202** | GCF_022398435.1 | PG_14 | 99.46 | 0.9 | Host-associated | Mollusca |
| **P_M8** | GCF_018141365.1 | PG_15 | 97.85 | 0.98 | Host-associated | Arthropoda |
| **P_maricaloris_LMG19692** | GCF_012641725.1 | PG_15 | 99.66 | 0.21 | Host-associated | Porifera |
| **P_MB47** | GCF_011319765.1 | PG_29 | 99.95 | 0.25 | Host-associated | Echinodermata |
| **P_OANN1** | GCF_024124685.1 | PG_15 | 99.75 | 0.18 | Host-associated | Cnidaria |
| **P_Of11M6** | GCF_022347905.1 | PG_15 | 99.73 | 0.73 | Host-associated | Cnidaria |
| **P_MEBiC03607** | GCF_004792295.1 | PG_26 | 99.95 | 0.72 | Free-living | Seawater |
| **P_OF5H5** | GCF_021530695.1 | PG_15 | 99.75 | 0.39 | Host-associated | Cnidaria |
| **P_MMG024** | GCF_021654175.1 | PG_53 | 99.75 | 0.63 | Host-associated | Annelida |
| **P_N12309** | GCF_032716425.1 | PG_25 | 99.95 | 0.18 | Free-living | Seawater |
| **P_NBRC102222** | GCF_007989585.1 | PG_46 | 99.49 | 0.51 | Free-living | Seawater |
| **P_NBRC12985** | GCF_006539245.1 | PG_47 | 99.84 | 0.25 | Free-living | Seawater |
| **P_neustronica_PAMC28425** | GCF_001653135.1 | PG_33 | 99.75 | 0.88 | Free-living | Seawater |
| **P_neustronica_SM1927** | GCF_007786355.1 | PG_32 | 99.75 | 1.03 | Free-living | Seawater |
| **P_NJ631** | GCF_000276645.1 | PG_15 | 100 | 0.16 | Host-associated | Porifera |
| **P_Of5H6** | GCF_040104565.1 | PG_15 | 99.75 | 0.39 | Host-associated | Cnidaria |
| **P_OF7H1** | GCF_022347825.1 | PG_15 | 100 | 0.53 | Host-associated | Cnidaria |
| **P_Of7M16** | GCF_022347805.1 | PG_01 | 99.70 | 0.51 | Host-associated | Cnidaria |
| **P_OFAV1** | GCF_021532595.1 | PG_26 | 99.66 | 3.78 | Host-associated | Cnidaria |
| **P_OOF1S7** | GCF_022347765.1 | PG_11 | 99.71 | 0.49 | Host-associated | Cnidaria |
| **P_bablabjr004** | GCF_016464215.1 | PG_26 | 99.70 | 0.74 | Host-associated | Cnidaria |
| **P_CnMc713** | GCF_040104495.1 | PG_16 | 99.37 | 0.72 | Host-associated | Cnidaria |
| **P_CnMc715** | GCF_022347925.1 | PG_16 | 99.37 | 0.72 | Host-associated | Cnidaria |
| **P_ostereae_hOe124** | GCF_029023665.1 | PG_31 | 99.12 | 1.26 | Host-associated | Mollusca |
| **P_ostereae_hOe125** | GCF_029026505.1 | PG_31 | 99.12 | 1.26 | Host-associated | Mollusca |
| **P_ostereae_hOe66** | GCF_018069805.1 | PG_31 | 99.12 | 1.26 | Host-associated | Mollusca |
| **P_P111** | GCF_001399995.1 | PG_45 | 100 | 0.81 | Host-associated | Cnidaria |
| **P_P18** | GCF_001399985.1 | PG_30 | 100 | 0.74 | Host-associated | Cnidaria |
| **P_P19** | GCF_001399975.1 | PG_53 | 99.75 | 0.82 | Host-associated | Cnidaria |
| **P_CnMc737** | GCF_024124485.1 | PG_30 | 99.95 | 1.01 | Host-associated | Cnidaria |
| **P_PA2MD11** | GCF_017808125.1 | PG_29 | 99.95 | 0.25 | Host-associated | Porifera |
| **P_McH142** | GCF_022347845.1 | PG_05 | 99.96 | 0.51 | Host-associated | Cnidaria |
| **P_peptidolytica_DSM14001** | GCF_012641745.1 | PG_13 | 95.32 | 1.22 | Free-living | Seawater |
| **P_peptidolytica_F1250A1** | GCF_014858745.1 | PG_13 | 99.49 | 1.22 | Free-living | Seawater |
| **P_peptidolytica_NBRC101021** | GCF_007989895.1 | PG_13 | 99.49 | 1.22 | Free-living | Seawater |
| **P_phenolica_KCTC12086** | GCF_001444405.1 | PG_49 | 99.95 | 1.12 | Free-living | Seawater |
| **P_phenolica_OBC302** | GCF_014925335.1 | PG_49 | 99.95 | 1.12 | Free-living | Seawater |
| **P_phenolica_S1093** | GCF_005886345.1 | PG_48 | 99.95 | 0.84 | Free-living | Seawater |
| **P_phenolica_S1189** | GCF_005887275.1 | PG_48 | 99.95 | 2.24 | Free-living | Seawater |
| **P_phenolica_S3663** | GCF_005876835.1 | PG_51 | 99.95 | 0.57 | Free-living | Seawater |
| **P_phenolica_S3898** | GCF_004214845.1 | PG_48 | 99.70 | 1.02 | Free-living | Seawater |
| **P_McH17** | GCF_013366255.1 | PG_13 | 99.75 | 1.47 | Host-associated | Cnidaria |
| **P_porphyrae_MNAD16** | GCF_003591235.1 | PG_28 | 99.95 | 0.53 | Free-living | Seawater |
| **P_PPB1** | GCF_015356275.1 | PG_05 | 99.01 | 0.25 | Host-associated | Porifera |
| **P_SMS1** | GCF_021530565.1 | PG_04 | 99.95 | 1.28 | Host-associated | Cnidaria |
| **P_XMcav11Q** | GCF_040104525.1 | PG_15 | 99.75 | 1.14 | Host-associated | Cnidaria |
| **P_rhizosphaerae_hCg42** | GCF_028885455.1 | PG_32 | 100 | 4.86 | Host-associated | Mollusca |
| **P_rubra_DSM6842** | GCF_000238295.3 | PG_07 | 99.71 | 0.99 | Free-living | Seawater |
| **P_rubra_S1946** | GCF_004212645.1 | PG_10 | 99.71 | 0.18 | Host-associated | Arthropoda |
| **P_rubra_S2471** | GCF_000967655.1 | PG_09 | 99.96 | 0.91 | Host-associated | Mollusca |
| **P_rubra_SCSIO_6842** | GCF_001482385.1 | PG_06 | 99.96 | 0.34 | Free-living | Seawater |
| **P_ruthenica_LMG19699** | GCF_008808095.1 | PG_16 | 98.85 | 1.47 | Host-associated | Mollusca |
| **P_ruthenica_S2756** | GCF_005876865.1 | PG_16 | 99.37 | 0.78 | Host-associated | Mollusca |
| **P_ruthenica_S3245** | GCF_005886975.1 | PG_16 | 99.49 | 2.52 | Host-associated | Arthropoda |
| **P_S2893** | GCF_005887105.1 | PG_45 | 100 | 0.79 | Host-associated | Echinodermata |
| **P_S3173** | GCF_005930795.1 | PG_34 | 99.67 | 0.76 | Host-associated | Cnidaria |
| **P_S3178** | GCF_005886985.1 | PG_47 | 99.96 | 2.25 | Host-associated | Arthropoda |
| **P_S3260** | GCF_005886925.1 | PG_34 | 99.95 | 1.49 | Host-associated | Arthropoda |
| **P_Scap03** | GCF_013393115.1 | PG_35 | 97.85 | 0.84 | Host-associated | Porifera |
| **P_scap25** | GCF_013394125.1 | PG_35 | 99.95 | 0.84 | Host-associated | Porifera |
| **P_scap26** | GCF_013394165.1 | PG_35 | 99.95 | 0.84 | Host-associated | Porifera |
| **P_SCSIO11900** | GCF_000576475.1 | PG_34 | 99.95 | 0.38 | Host-associated | Cnidaria |
| **P_XMcav2N** | GCF_024124545.1 | PG_08 | 99.92 | 0.38 | Host-associated | Cnidaria |
| **P_XMcav2N2** | GCF_040104535.1 | PG_08 | 99.92 | 0.38 | Host-associated | Cnidaria |
| **P_2102** | GCF_013350045.1 | PG_30 | 99.95 | 0.44 | Host-associated | Cnidaria |
| **P_2103** | GCF_013349985.1 | PG_30 | 99.95 | 0.44 | Host-associated | Cnidaria |
| **P_SDCH90** | GCF_019134595.1 | PG_30 | 99.44 | 0.99 | Host-associated | Mollusca |
| **P_SG411** | GCF_014164575.1 | PG_32 | 99.75 | 1.05 | Free-living | Seawater |
| **P_SG412** | GCF_014164535.1 | PG_32 | 99.87 | 1.03 | Free-living | Seawater |
| **P_SG415** | GCF_014164495.1 | PG_32 | 98.97 | 1.72 | Free-living | Seawater |
| **P_SG416** | GCF_014164455.1 | PG_38 | 99.73 | 0.41 | Free-living | Seawater |
| **P_SG418** | GCF_014164425.1 | PG_32 | 99.62 | 1.53 | Free-living | Seawater |
| **P_SG431** | GCF_014164415.1 | PG_40 | 99.94 | 0.37 | Free-living | Seawater |
| **P_SG433** | GCF_014164385.1 | PG_40 | 100 | 0.21 | Free-living | Seawater |
| **P_SG434** | GCF_014164325.1 | PG_40 | 99.75 | 0.22 | Free-living | Seawater |
| **P_SG435** | GCF_014164345.1 | PG_40 | 100 | 0.13 | Free-living | Seawater |
| **P_SG436** | GCF_014164365.1 | PG_32 | 99.87 | 1.56 | Free-living | Seawater |
| **P_SG437** | GCF_014164295.1 | PG_32 | 99.37 | 1.31 | Free-living | Seawater |
| **P_SG438** | GCF_014164245.1 | PG_40 | 99.92 | 0.16 | Free-living | Seawater |
| **P_SG441** | GCF_014164315.1 | PG_32 | 99.87 | 1.65 | Free-living | Seawater |
| **P_SG4417** | GCF_014164235.1 | PG_32 | 99.96 | 0.86 | Free-living | Seawater |
| **P_SG444** | GCF_014164225.1 | PG_38 | 100 | 0.38 | Free-living | Seawater |
| **P_SG445** | GCF_014164175.1 | PG_37 | 99.97 | 0.32 | Free-living | Seawater |
| **P_SG448** | GCF_014164125.1 | PG_32 | 100 | 0.74 | Free-living | Seawater |
| **P_SG451** | GCF_014164135.1 | PG_40 | 100 | 0.41 | Free-living | Seawater |
| **P_SG452** | GCF_014164115.1 | PG_40 | 100 | 0.13 | Free-living | Seawater |
| **P_SG453** | GCF_014164075.1 | PG_40 | 100 | 0.21 | Free-living | Seawater |
| **P_SG455** | GCF_014164085.1 | PG_38 | 100 | 0.13 | Free-living | Seawater |
| **P_SG456** | GCF_014164025.1 | PG_40 | 99.66 | 0.71 | Free-living | Seawater |
| **P_303** | GCF_013349995.1 | PG_30 | 99.95 | 0.72 | Host-associated | Cnidaria |
| **P_802** | GCF_013350085.1 | PG_30 | 99.95 | 0.44 | Host-associated | Cnidaria |
| **P_shioyasakiensis_BMC2** | GCF_029992275.1 | PG_30 | 99.95 | 0.44 | Host-associated | Cnidaria |
| **P_shioyasakiensis_BMC3** | GCF_029992495.1 | PG_30 | 99.89 | 0.44 | Host-associated | Cnidaria |
| **P_shioyasakiensis_BMC4** | GCF_029991435.1 | PG_30 | 99.95 | 0.80 | Host-associated | Cnidaria |
| **P_shioyasakiensis_G21653S1** | GCF_027213805.1 | PG_30 | 99.95 | 0.79 | Host-associated | Cnidaria |
| **P_shioyasakiensis_LC2** | GCF_030247135.1 | PG_30 | 99.95 | 1.36 | Free-living | Seawater |
| **P_shioyasakiensis_M1400201** | GCF_022398475.1 | PG_30 | 99.49 | 0.68 | Host-associated | Mollusca |
| **P_SiA1** | GCF_018732205.1 | PG_34 | 99.44 | 0.51 | Host-associated | Echinodermata |
| **P_SK18** | GCF_001974855.1 | PG_36 | 99.70 | 0.51 | Host-associated | Porifera |
| **P_SK20** | GCF_001974845.1 | PG_34 | 99.92 | 0.74 | Host-associated | Porifera |
| **P_shioyasakiensis_BMC5** | GCF_029991425.1 | PG_30 | 99.95 | 0.44 | Host-associated | Cnidaria |
| **P_spongiae_SAO44** | GCF_002814155.1 | PG_52 | 100 | 2.17 | Free-living | Seawater |
| **P_spongiae_UST010723006** | GCF_000238255.3 | PG_52 | 100 | 0.38 | Host-associated | Porifera |
| **P_SR411** | GCF_014164015.1 | PG_40 | 100 | 0.16 | Free-living | Seawater |
| **P_SR414** | GCF_014163985.1 | PG_32 | 99.75 | 0.83 | Free-living | Seawater |
| **P_SR415** | GCF_014163975.1 | PG_40 | 100 | 0.16 | Free-living | Seawater |
| **P_SR416** | GCF_014163945.1 | PG_32 | 99.62 | 1.56 | Free-living | Seawater |
| **P_SR417** | GCF_014163935.1 | PG_40 | 100 | 0.24 | Free-living | Seawater |
| **P_SR418** | GCF_014163915.1 | PG_32 | 100 | 0.92 | Free-living | Seawater |
| **P_SR432** | GCF_014163885.1 | PG_40 | 100 | 0.21 | Free-living | Seawater |
| **P_SR433** | GCF_014163875.1 | PG_40 | 99.24 | 0.13 | Free-living | Seawater |
| **P_SR435** | GCF_014163835.1 | PG_40 | 100 | 0.16 | Free-living | Seawater |
| **P_SR436** | GCF_014163815.1 | PG_40 | 100 | 0.16 | Free-living | Seawater |
| **P_SR437** | GCF_014163785.1 | PG_40 | 100 | 0.57 | Free-living | Seawater |
| **P_SR442** | GCF_014163745.1 | PG_40 | 99.87 | 0.16 | Free-living | Seawater |
| **P_SR445** | GCF_014163675.1 | PG_32 | 99.62 | 1.56 | Free-living | Seawater |
| **P_SR448** | GCF_014163635.1 | PG_32 | 100 | 0.96 | Free-living | Seawater |
| **P_SR451** | GCF_014163655.1 | PG_40 | 100 | 0.47 | Free-living | Seawater |
| **P_SR454** | GCF_014163535.1 | PG_37 | 99.72 | 1.06 | Free-living | Seawater |
| **P_SR455** | GCF_014163565.1 | PG_40 | 96.96 | 1.61 | Free-living | Seawater |
| **P_SR456** | GCF_014163545.1 | PG_32 | 99.62 | 1.65 | Free-living | Seawater |
| **P_SW010604** | GCF_001293805.1 | PG_16 | 99.12 | 1.03 | Free-living | Seawater |
| **P_TB25** | GCF_000497995.1 | PG_40 | 95.32 | 0.73 | Host-associated | Porifera |
| **P_tetradonis_CSB01KR** | GCF_001723425.1 | PG_34 | 99.95 | 0.13 | Host-associated | Echinodermata |
| **P_translucida_KMM520** | GCF_001465295.1 | PG_37 | 100 | 0.04 | Free-living | Seawater |
| **P_tunicata_D2** | GCF_002310815.1 | PG_56 | 99.75 | 01.3 | Host-associated | Chordata |
| **P_UG31** | GCF_037120685.1 | PG_12 | 99.66 | 0.85 | Host-associated | Arthropoda |
| **P_UG32** | GCF_037120705.1 | PG_12 | 100 | 0.18 | Host-associated | Arthropoda |
| **P_undina_DSM6065** | GCF_000238275.3 | PG_35 | 99.87 | 0.25 | Free-living | Seawater |
| **P_undina_NEM01** | GCF_048401115.1 | PG_35 | 100 | 0.71 | Host-associated | Nematoda |
| **P_undina_NEM02** | GCF_048401065.1 | PG_35 | 100 | 0.71 | Host-associated | Nematoda |
| **P_viridis_BBR56** | GCF_017742995.1 | PG_10 | 99.20 | 0.51 | Free-living | Seawater |
| **P_xiamenensis_PSDB** | GCF_030994125.1 | PG_23 | 98.08 | 2.31 | Host-associated | Mollusca |
| **P_bablabjr010** | GCF_016464105.1 | PG_29 | 99.95 | 0.75 | Host-associated | Cnidaria |
| **P_PAST1** | GCF_021532515.1 | PG_29 | 99.49 | 1.01 | Host-associated | Cnidaria |
| **P_SCSIO43095** | GCF_023650455.1 | PG_34 | 99.02 | 0.63 | Host-associated | Cnidaria |
