## Supplementary Table 2 for "*Pseudoalteromonas* is a novel symbiont of marine invertebrates that exhibits broad patterns of phylosymbiosis"

**Supplementary Table 2: Summary of single-worm metagenomic samples used to identify the prevalence of *Pseudoalteromonas* in the microbiome of marine nematodes.** Metadata includes the habitat that the worms were isolated from, the genus-level identification of the worm, the genome amplification protocol, and the date they were collected from.

| **Nematode ID** | **SampleID** | **Year Collected** | **Month Collected** | **Habitat** | **Genus** | **Amplification** | **Number of Worms** | **Seq Facility** |
| --- | --- | --- | --- | --- | --- | --- | --- | --- |
| **enoplolaimus.1** | enopolaimus_001_AQ_202010 | 2020 | October | Antarctica | Enopolaimus | No | 1 | JGI |
| **enoplolaimus.2** | enopolaimus_002_AQ_202010 | 2020 | October | Antarctica | Enopolaimus | No | 1 | JGI |
| **pontonema.1** | pontonema_016_AQ_202010 | 2020 | October | Antarctica | Pontonema | No | 1 | JGI |
| **pontonema.2** | pontonema_015_AQ_202010 | 2020 | October | Antarctica | Pontonema | No | 1 | JGI |
| **pontonema.3** | pontonema_017_AQ_202010 | 2020 | October | Antarctica | Pontonema | No | 1 | JGI |
| **thoracostoma.1** | thoracostoma_021_AQ_202010 | 2020 | October | Antarctica | Thoracostoma | No | 1 | JGI |
| **thoracostoma.2** | thoracostoma_023_AQ_202010 | 2018 | July | Antarctica | Thoracostoma | No | 1 | JGI |
| **thoracostoma.3** | thoracostoma_025_AQ_202010 | 2020 | October | Antarctica | Thoracostoma | No | 1 | JGI |
| **thoracostoma.4** | thoracostoma_026_AQ_202010 | 2020 | October | Antarctica | Thoracostoma | No | 1 | JGI |
| **thoracostoma.5** | thoracostoma_027_AQ_202010 | 2020 | October | Antarctica | Thoracostoma | No | 1 | JGI |
| **thoracostoma.6** | thoracostoma_028_AQ_202010 | 2020 | October | Antarctica | Thoracostoma | No | 1 | JGI |
| **thoracostoma.7** | thoracostoma_029_AQ_202010 | 2020 | October | Antarctica | Thoracostoma | No | 1 | JGI |
| **thoracostoma.8** | thoracostoma_030_AQ_202010 | 2020 | October | Antarctica | Thoracostoma | No | 1 | JGI |
| **thoracostoma.9** | thoracostoma_031_AQ_202010 | 2020 | October | Antarctica | Thoracostoma | No | 1 | JGI |
| **thoracostoma.10** | thoracostoma_022_AQ_201807 | 2018 | July | Bodega Bay | Thoracostoma | No | 1 | JGI |
| **thoracostoma.11** | thoracostoma_024_AQ_201807 | 2018 | July | Bodega Bay | Thoracostoma | No | 1 | JGI |
| **oncholaimellus.1** | oncholaimellus_94_DR_FL_202212 | 2022 | December | Dolphins Reef | Oncholaimellus | No | 1 | JGI |
| **pareurystomina.1** | pareurstomina_93_DR_FL_202212 | 2022 | December | Dolphins Reef | Pareurystomina | No | 1 | JGI |
| **oncholaimidae.1** | oncholaimidae_006_GA_TI_202202 | 2022 | December | Tybee Island | Oncholaimidae | No | 1 | JGI |
| **oncholaimidae.2** | oncholaimidae_007_GA_TI_202202 | 2022 | December | Tybee Island | Oncholaimidae | No | 1 | JGI |
| **adoncholaimus.1** | adoncholaimus_032_GA_TI_202107 | 2021 | July | Tybee Island | Adoncholaimus | GenomiPhi | 1 | UGA |
| **adoncholaimus.10** | adoncholaimus_041_GA_TI_202202 | 2022 | February | Tybee Island | Adoncholaimus | No | 3 | UGA |
| **adoncholaimus.11** | adoncholaimus_042_GA_TI_202202 | 2022 | February | Tybee Island | Adoncholaimus | No | 5 | UGA |
| **adoncholaimus.2** | adoncholaimus_033_GA_TI_202107 | 2021 | July | Tybee Island | Adoncholaimus | GenomiPhi | 1 | UGA |
| **adoncholaimus.3** | adoncholaimus_034_GA_TI_202107 | 2021 | July | Tybee Island | Adoncholaimus | GenomiPhi | 1 | UGA |
| **adoncholaimus.4** | adoncholaimus_035_GA_TI_202107 | 2021 | July | Tybee Island | Adoncholaimus | No | 1 | UGA |
| **adoncholaimus.5** | adoncholaimus_036_GA_TI_202107 | 2021 | July | Tybee Island | Adoncholaimus | No | 1 | UGA |
| **adoncholaimus.6** | adoncholaimus_037_GA_TI_202107 | 2021 | July | Tybee Island | Adoncholaimus | GenomiPhi | 1 | UGA |
| **adoncholaimus.7** | adoncholaimus_038_GA_TI_202202 | 2022 | February | Tybee Island | Adoncholaimus | GenomiPhi | 1 | UGA |
| **adoncholaimus.8** | adoncholaimus_039_GA_TI_202202 | 2022 | February | Tybee Island | Adoncholaimus | No | 10 | UGA |
| **adoncholaimus.9** | adoncholaimus_040_GA_TI_202202 | 2022 | February | Tybee Island | Adoncholaimus | No | 20 | UGA |
| **epacanthion.1** | epacanthion_003_GA_TI_202311 | 2023 | November | Tybee Island | Epacanthion | Yes | 1 | JGI |
| **epacanthion.2** | epacanthion_004_GA_TI_202311 | 2023 | November | Tybee Island | Epacanthion | Yes | 1 | JGI |
| **epacanthion.3** | epacanthion_005_GA_TI_202311 | 2023 | November | Tybee Island | Epacanthion | Yes | 1 | JGI |
| **metoncholaimus.1** | metoncholaimus_043_201807 | 2018 | July | Tybee Island | Metoncholaimus | No | 1 | JGI |
| **oncholaimidae.3** | oncholaimidae_008_GA_TI_202311 | 2023 | November | Tybee Island | Oncholaimidae | Yes | 1 | JGI |
| **oncholaimidae.4** | oncholaimidae_009_GA_TI_202311 | 2023 | November | Tybee Island | Oncholaimidae | Yes | 1 | JGI |
| **oncholaimidae.5** | oncholaimidae_010_GA_TI_202311 | 2023 | November | Tybee Island | Oncholaimidae | Yes | 1 | JGI |
| **oncholaimidae.6** | oncholaimidae_011_GA_TI_202311 | 2023 | November | Tybee Island | Oncholaimidae | Yes | 1 | JGI |
| **oncholaimidae.7** | oncholaimidae_012_GA_TI_202311 | 2023 | November | Tybee Island | Oncholaimidae | Yes | 1 | JGI |
| **oncholaimidae.8** | oncholaimidae_013_GA_TI_202311 | 2023 | November | Tybee Island | Oncholaimidae | Yes | 1 | JGI |
| **oncholaimidae.9** | oncholaimidae_014_GA_TI_202311 | 2023 | November | Tybee Island | Oncholaimidae | Yes | 1 | JGI |
| **sphaerolaimus.1** | sphaerolaimus_019_GA_TI_202202 | 2022 | February | Tybee Island | Sphaerolaimus | No | 1 | JGI |
| **sphaerolaimus.2** | sphaerolaimus_018_GA_TI_202202 | 2022 | February | Tybee Island | Sphaerolaimus | No | 1 | JGI |
| **sphaerolaimus.3** | sphaerolaimus_020_GA_TI_202311 | 2023 | November | Tybee Island | Sphaerolaimus | Yes | 1 | JGI |
