## Supplementary Table 3 for "*Pseudoalteromonas* is a novel symbiont of marine invertebrates that exhibits broad patterns of phylosymbiosis"

**Supplementary Table 3: Accessory genes more commonly found in host-associated genomes.** A summary of the genes more commonly found in host-associated (t+) pigmented (orange) and nonpigmented (purple) genomes**.** The gene family ID is in bold, followed by a brief description of the product. The presence and absence of a function or gene is indicated by (g+) or (g-), respectively. Functional predictions of hypothetical proteins by Gaia, an AI agent developed by Tatta Bio that integrates several genomic and structural features to predict functions of novel proteins, are in red. An odds-ratio greater than one indicates that the gene/function is more commonly associated with the host-associated lifestyle strains**.**

| **Gene** | **g+t+** | **g+t-** | **g-t+** | **g-t-** | **sensitivity** | **specificity** | **Odds-Ratio** | **Fisher P** | **Lifestyle** | **Clade** |
| --- | --- | --- | --- | --- | --- | --- | --- | --- | --- | --- |
| **g001815_1_000001** (DmlR)  HTH-type transcriptional regulator | 56 | 9 | 10 | 13 | 84.85 | 59.09 | 8.09 | 0.0001 | Host-associated | Pigmented |
| **g000097_000001** (AcuC)  Acetoin utilization protein | 64 | 13 | 2 | 9 | 96.97 | 40.91 | 22.15 | 3.47E-05 | Host-associated | Pigmented |
| **g004459**  periplasmic substrate-binding protein | 36 | 2 | 30 | 20 | 54.55 | 90.91 | 12.00 | 0.0001 | Host-associated | Pigmented |
| **g002208_1_000003** (YdhP)  Inner membrane transport protein | 55 | 8 | 11 | 14 | 83.33 | 63.64 | 8.75 | 6.34E-05 | Host-associated | Pigmented |
| **g001387_1_000001**  transport of complex carbohydrates | 58 | 10 | 8 | 12 | 87.88 | 54.55 | 8.70 | 0.0001 | Host-associated | Pigmented |
| **g001123**  bacteriocin immunity or production | 43 | 4 | 23 | 18 | 65.15 | 81.82 | 8.413 | 0.0001 | Host-associated | Pigmented |
| **g012839** (exbD)  Biopolymer transport protein | 13 | 0 | 65 | 69 | 16.67 | 100.00 | inf | 0.0002 | Host-associated | Nonpigmented |
| **g004033_1_000001**  DNA transposase | 13 | 0 | 65 | 69 | 16.67 | 100.00 | inf | 0.0002 | Host-associated | Nonpigmented |
| **g004526_1_000001**  protein toxin complex | 15 | 0 | 63 | 69 | 19.23 | 100.00 | inf | 4.18E-05 | Host-associated | Nonpigmented |
| **g005471_000003**  membrane-targeting toxin protein | 13 | 0 | 65 | 69 | 16.67 | 100.00 | inf | 0.0002 | Host-associated | Nonpigmented |
| **g006486_000001**  transcriptional regulator | 17 | 0 | 61 | 69 | 21.79 | 100.00 | inf | 9.15E-06 | Host-associated | Nonpigmented |
| **g009813**  antimicrobial peptide precursor | 16 | 0 | 62 | 69 | 20.51 | 100.00 | inf | 1.96E-05 | Host-associated | Nonpigmented |
| **g007355** (irtB)  iron import permease protein | 14 | 0 | 64 | 69 | 17.95 | 100.00 | inf | 8.85E-05 | Host-associated | Nonpigmented |
| **g000623_000001**  membrane metal-binding protein | 77 | 44 | 1 | 25 | 98.72 | 36.23 | 43.75 | 6.25E-09 | Host-associated | Nonpigmented |
| **g009670_000003** (higA-1)  Antitoxin HigA-1 | 19 | 1 | 59 | 68 | 24.36 | 98.55 | 21.90 | 2.21E-05 | Host-associated | Nonpigmented |
| **g008227** (pgrR)  HTH-type transcriptional regulator | 17 | 1 | 61 | 68 | 21.79 | 98.55 | 18.95 | 9.26E-05 | Host-associated | Nonpigmented |
| **g005408_000002**  oxygen-responsive signaling system | 17 | 1 | 61 | 68 | 21.79 | 98.55 | 18.95 | 9.26E-05 | Host-associated | Nonpigmented |
| **g009294**  phosphate starvation-induced protein | 16 | 1 | 62 | 68 | 20.51 | 98.55 | 17.55 | 0.0002 | Host-associated | Nonpigmented |
| **g003698_1_000001** (mbtH)  nonribosomal peptide protein | 25 | 2 | 53 | 67 | 32.05 | 97.10 | 15.80 | 2.59E-06 | Host-associated | Nonpigmented |
| **g000192_000002** (ygeA)  L-aspartate/glutamate racemase | 69 | 25 | 9 | 44 | 88.46 | 63.77 | 13.49 | 4.02E-11 | Host-associated | Nonpigmented |
| **g002157_000001**  metal ion or porphyrin transporter | 74 | 43 | 4 | 26 | 94.87 | 37.68 | 11.19 | 8.50E-07 | Host-associated | Nonpigmented |
| **g004055_000001** (hchA)  Protein/nucleic acid deglycase | 34 | 5 | 44 | 64 | 43.59 | 92.75 | 9.89 | 4.99E-07 | Host-associated | Nonpigmented |
| **g001928**  periplasmic oxidoreductase | 58 | 16 | 20 | 53 | 74.36 | 76.81 | 9.61 | 4.66E-10 | Host-associated | Nonpigmented |
| **g003177**  acylphosphatase | 37 | 6 | 41 | 63 | 47.44 | 91.30 | 9.48 | 1.71E-07 | Host-associated | Nonpigmented |
| **g005155**  transport or regulatory system | 23 | 3 | 55 | 66 | 29.49 | 95.65 | 9.20 | 5.56E-05 | Host-associated | Nonpigmented |
| **g004624_000001** (fabL)  Enoyl-[acyl-carrier-protein] reductase | 28 | 4 | 50 | 65 | 35.90 | 94.20 | 9.10 | 6.40E-06 | Host-associated | Nonpigmented |
| **g003915**  N-acetyltransferase | 22 | 3 | 56 | 66 | 28.21 | 95.65 | 8.64 | 0.0001 | Host-associated | Nonpigmented |
| **g004578** (tolB)  Tol-Pal system protein | 22 | 3 | 56 | 66 | 28.21 | 95.65 | 8.64 | 0.0001 | Host-associated | Nonpigmented |
| **g002150_1_000001**  protein dephosphorylation | 31 | 5 | 47 | 64 | 39.74 | 92.75 | 8.44 | 2.90E-06 | Host-associated | Nonpigmented |
| **g003352_000001**  membrane iron transporter | 54 | 15 | 24 | 54 | 69.23 | 78.26 | 8.10 | 9.63E-09 | Host-associated | Nonpigmented |
| **g002367_000003** (abgT)  p-aminobenzoyl-glutamate transport | 74 | 48 | 4 | 21 | 94.87 | 30.43 | 8.09 | 4.61E-05 | Host-associated | Nonpigmented |
| **g004521_1_000005** (ywrD)  glutathione hydrolase | 45 | 10 | 33 | 59 | 57.69 | 85.51 | 8.05 | 6.41E-08 | Host-associated | Nonpigmented |
| **g002861_000002**  phosphodiesterase/phospholipase antimicrobial system | 32 | 6 | 46 | 63 | 41.03 | 91.30 | 7.30 | 8.45E-06 | Host-associated | Nonpigmented |
| **g006714** (pupA)  Ferric-pseudobactin receptor | 24 | 4 | 54 | 65 | 30.77 | 94.20 | 7.22 | 0.0001 | Host-associated | Nonpigmented |
| **g003015 (hchA)**  protein/nucleic acid deglycase | 28 | 5 | 50 | 64 | 35.90 | 92.75 | 7.17 | 2.43E-05 | Host-associated | Nonpigmented |
| **g002070_000001**  N-acetyltransferase | 50 | 14 | 28 | 55 | 64.10 | 79.71 | 7.02 | 7.54E-08 | Host-associated | Nonpigmented |
| **g000120_1_000001** (aprE)  extracellular protease | 63 | 26 | 15 | 43 | 80.77 | 62.32 | 6.95 | 1.04E-07 | Host-associated | Nonpigmented |
| **g000121_2_000002** (katG2)  Catalase-peroxidase | 70 | 39 | 8 | 30 | 89.74 | 43.48 | 6.73 | 4.35E-06 | Host-associated | Nonpigmented |
| **g005284** (ybhF)  ATP-binding multidrug transporter | 39 | 9 | 39 | 60 | 50.00 | 86.96 | 6.67 | 1.49E-06 | Host-associated | Nonpigmented |
| **g005541** (ybhS)  multidrug transporter permease | 39 | 9 | 39 | 60 | 50.00 | 86.96 | 6.67 | 1.49E-06 | Host-associated | Nonpigmented |
| **g003527** (cobC)  Adenosylcobalamin/alpha-ribazole phosphatase | 26 | 5 | 52 | 64 | 33.33 | 92.75 | 6.40 | 9.36E-05 | Host-associated | Nonpigmented |
| **g005334** (YciF)  osmotic stress response system | 26 | 5 | 52 | 64 | 33.33 | 92.75 | 6.40 | 9.36E-05 | Host-associated | Nonpigmented |
| **g004086** (rbn)  ribonuclease | 26 | 5 | 52 | 64 | 33.33 | 92.75 | 6.40 | 9.36E-05 | Host-associated | Nonpigmented |
| **g001099_000002**  glycosyl hydrolase | 70 | 40 | 8 | 29 | 89.74 | 42.03 | 6.34 | 9.48E-06 | Host-associated | Nonpigmented |
| **g005310** (mntP)  manganese efflux pump | 38 | 9 | 40 | 60 | 48.72 | 86.96 | 6.33 | 3.13E-06 | Host-associated | Nonpigmented |
| **g002777_000002**  secondary transporter for gluconate | 61 | 25 | 17 | 44 | 78.21 | 63.77 | 6.32 | 3.27E-07 | Host-associated | Nonpigmented |
| **g004106_000001** (bdhA)  D-beta-hydroxybutyrate dehydrogenase | 51 | 16 | 27 | 53 | 65.38 | 76.81 | 6.26 | 4.36E-07 | Host-associated | Nonpigmented |
| **g003882_000001** (metXA)  homoserine O-acetyltransferase | 51 | 16 | 27 | 53 | 65.38 | 76.81 | 6.26 | 4.36E-07 | Host-associated | Nonpigmented |
| **g004139_000002** (fyuA)  pesticin receptor | 51 | 16 | 27 | 53 | 65.38 | 76.81 | 6.26 | 4.36E-07 | Host-associated | Nonpigmented |
| **g005346_000001** (DegU)  transcriptional regulator | 51 | 16 | 27 | 53 | 65.38 | 76.81 | 6.26 | 4.36E-07 | Host-associated | Nonpigmented |
| **g003571_000002**  cell surface protein involved in cell adhesion or biofilm formation | 40 | 10 | 38 | 59 | 51.28 | 85.51 | 6.21 | 2.14E-06 | Host-associated | Nonpigmented |
| **g001613** (LPtD)  surface carbohydrate-binding protein | 57 | 21 | 21 | 48 | 73.08 | 69.57 | 6.20 | 2.38E-07 | Host-associated | Nonpigmented |
| **g001903** (scrK)  fructokinase | 46 | 13 | 32 | 56 | 58.97 | 81.16 | 6.19 | 7.41E-07 | Host-associated | Nonpigmented |
| **g003572_000001** (rcsC)  sensor histidine kinase | 52 | 17 | 26 | 52 | 66.67 | 75.36 | 6.12 | 4.71E-07 | Host-associated | Nonpigmented |
| **gr_000001**  phospholipid biosynthesis | 25 | 5 | 53 | 64 | 32.05 | 92.75 | 6.04 | 0.0002 | Host-associated | Nonpigmented |

| **g004229_000001**  iron-dependent dioxygenase | 25 | 5 | 53 | 64 | 32.05 | 92.75 | 6.04 | 0.0002 | Host-associated | Nonpigmented |
| --- | --- | --- | --- | --- | --- | --- | --- | --- | --- | --- |
| **g001883_000001** (allB)  allantoinase | 45 | 13 | 33 | 56 | 57.69 | 81.16 | 5.87 | 1.64E-06 | Host-associated | Nonpigmented |
| **g001456**  membrane organization or lipid metabolism | 57 | 22 | 21 | 47 | 73.08 | 68.12 | 5.80 | 5.93E-07 | Host-associated | Nonpigmented |
| **g004021_1_000001** (sasA)  adaptive-response sensory kinase | 36 | 9 | 42 | 60 | 46.15 | 86.96 | 5.71 | 1.32E-05 | Host-associated | Nonpigmented |
| **g001949** (btuB)  vitamin B12 transporter | 62 | 28 | 16 | 41 | 79.49 | 59.42 | 5.67 | 1.60E-06 | Host-associated | Nonpigmented |
| **g000323_1_000003** (chiA)  chitinase | 67 | 36 | 11 | 33 | 85.90 | 47.83 | 5.58 | 1.08E-05 | Host-associated | Nonpigmented |
| **g002068_000001**  spheroidene monooxygenase; bleomycin resistance | 44 | 13 | 34 | 56 | 56.41 | 81.16 | 5.57 | 3.54E-06 | Host-associated | Nonpigmented |
| **g004062**  amino acid membrane transporter | 27 | 6 | 51 | 63 | 34.62 | 91.30 | 5.56 | 0.0002 | Host-associated | Nonpigmented |
| **g_000001**  regulates iron acquisition | 27 | 6 | 51 | 63 | 34.62 | 91.30 | 5.56 | 0.0002 | Host-associated | Nonpigmented |
| **g004159** (nfdA)  N-substitute formamide deformylase | 27 | 6 | 51 | 63 | 34.62 | 91.30 | 5.56 | 0.0002 | Host-associated | Nonpigmented |
| **g000761_000001**  linoleate hydratase | 69 | 40 | 9 | 29 | 88.46 | 42.03 | 5.56 | 2.74E-05 | Host-associated | Nonpigmented |
| **g_000001**  arginine/agmatine metabolism | 68 | 38 | 10 | 31 | 87.18 | 44.93 | 5.55 | 1.73E-05 | Host-associated | Nonpigmented |
| **g004027** (sasA)  adaptive-response sensory-kinase | 30 | 7 | 48 | 62 | 38.46 | 89.86 | 5.54 | 0.0001 | Host-associated | Nonpigmented |
| **g004173**  efflux transport system involved in multidrug resistance | 30 | 7 | 48 | 62 | 38.46 | 89.86 | 5.54 | 0.0001 | Host-associated | Nonpigmented |
| **g000943_1_000001**  holothin acyltransferase | 50 | 17 | 28 | 52 | 64.10 | 75.36 | 5.46 | 2.53E-06 | Host-associated | Nonpigmented |
| **g001232_000001**  metalloenzyme involved in cell wall component modification | 50 | 17 | 28 | 52 | 64.10 | 75.36 | 5.46 | 2.53E-06 | Host-associated | Nonpigmented |
| **g002926_000001** (yqjF)  inner membrane protein | 47 | 15 | 31 | 54 | 60.26 | 78.26 | 5.46 | 2.41E-06 | Host-associated | Nonpigmented |
| **g000056_2_000004** (dpp5)  dipeptidyl-peptidase | 56 | 22 | 22 | 47 | 71.79 | 68.12 | 5.44 | 1.41E-06 | Host-associated | Nonpigmented |
| **g003452_000003** (cstA)  peptide transporter | 57 | 23 | 21 | 46 | 73.08 | 66.67 | 5.43 | 1.43E-06 | Host-associated | Nonpigmented |
| **g003453_1_000001** (eptA)  Phosphoethanolamine transferase | 35 | 9 | 43 | 60 | 44.87 | 86.96 | 5.43 | 2.64E-05 | Host-associated | Nonpigmented |
| **g001094_000001** (czcA)  cobalt-zinc-cadmium resistance protein | 71 | 45 | 7 | 24 | 91.03 | 34.78 | 5.41 | 0.0002 | Host-associated | Nonpigmented |
| **g005980_000001** (hcaR)  hca transcriptional activator | 37 | 10 | 41 | 59 | 47.44 | 85.51 | 5.32 | 1.82E-05 | Host-associated | Nonpigmented |
| **g003060**  serine acetyltransferase | 37 | 10 | 41 | 59 | 47.44 | 85.51 | 5.32 | 1.82E-05 | Host-associated | Nonpigmented |
| **g002620**  membrane protein involved in cell organization or protein secretion | 32 | 8 | 46 | 61 | 41.03 | 88.41 | 5.30 | 7.38E-05 | Host-associated | Nonpigmented |
| **g001556** (degA)  HTH-type transcriptional regulator | 70 | 43 | 8 | 26 | 89.74 | 37.68 | 5.29 | 0.0001 | Host-associated | Nonpigmented |
| **g003693_000001**  penicillinase repressor that regulates antibiotic resistance genes | 39 | 11 | 39 | 58 | 50.00 | 84.06 | 5.27 | 1.23E-05 | Host-associated | Nonpigmented |
| **g003826**  membrane-associated stress response system | 29 | 7 | 49 | 62 | 37.18 | 89.86 | 5.24 | 0.0002 | Host-associated | Nonpigmented |
| **g001867_000001** (pepA)  Cytosol aminopeptidase | 69 | 41 | 9 | 28 | 88.46 | 40.58 | 5.24 | 5.63E-05 | Host-associated | Nonpigmented |
| **g000972** (btuB)  vitamin B12 transporter | 68 | 39 | 10 | 30 | 87.18 | 43.48 | 5.23 | 3.59E-05 | Host-associated | Nonpigmented |
| **g002796**  phosphate-specific outer membrane protein | 49 | 17 | 29 | 52 | 62.82 | 75.36 | 5.17 | 3.32E-06 | Host-associated | Nonpigmented |
| **g004891** (malT)  HTH-type transcriptional regulator | 34 | 9 | 44 | 60 | 43.59 | 86.96 | 5.15 | 5.19E-05 | Host-associated | Nonpigmented |
| **g003842**  curlin subunit that contributes to biofilm formation and bacterial surface adhesion | 34 | 9 | 44 | 60 | 43.59 | 86.96 | 5.15 | 5.19E-05 | Host-associated | Nonpigmented |
| **g005382_000001** (hcnA)  hydrogen cyanide synthase subunit | 31 | 8 | 47 | 61 | 39.74 | 88.41 | 5.03 | 0.0001 | Host-associated | Nonpigmented |
| **g003864_000001** (hcnB)  hydrogen cyanide synthase subunit | 31 | 8 | 47 | 61 | 39.74 | 88.41 | 5.03 | 0.0001 | Host-associated | Nonpigmented |
| **g004742_000003** (hcnC)  hydrogen cyanide synthase subunit | 31 | 8 | 47 | 61 | 39.74 | 88.41 | 5.03 | 0.0001 | Host-associated | Nonpigmented |
| **g001192**  secreted hydrolase/esterase enzyme | 47 | 16 | 31 | 53 | 60.26 | 76.81 | 5.02 | 6.04E-06 | Host-associated | Nonpigmented |
| **g001168_000001** (ylhS)  sulfoquinovose isomerase | 51 | 19 | 27 | 50 | 65.38 | 72.46 | 4.97 | 6.35E-06 | Host-associated | Nonpigmented |
| **g001112**  conjugated linoleic acid biosynthesis isomerase | 68 | 40 | 10 | 29 | 87.18 | 42.03 | 4.93 | 7.29E-05 | Host-associated | Nonpigmented |
| **g001708** (fucP)  L-fucose-proton symporter | 69 | 42 | 9 | 27 | 88.46 | 39.13 | 4.93 | 0.0001 | Host-associated | Nonpigmented |
| **g002520**  periplasmic substrate-binding protein involved in transport of polar amino acids | 33 | 9 | 45 | 60 | 42.31 | 86.96 | 4.89 | 0.0001 | Host-associated | Nonpigmented |
| **g002059_1_000001**  metallo-hydrolase | 53 | 21 | 25 | 48 | 67.95 | 69.57 | 4.85 | 6.95E-06 | Host-associated | Nonpigmented |
| **g002749_000001** (dnaE2)  error-prone DNA polymerase | 35 | 10 | 43 | 59 | 44.87 | 85.51 | 4.80 | 6.96E-05 | Host-associated | Nonpigmented |
| **g000732_1_000001**  periplasmic sugar-binding protein | 55 | 23 | 23 | 46 | 70.51 | 66.67 | 4.78 | 7.36E-06 | Host-associated | Nonpigmented |
| **g000017_2** (btuB)  vitamin B12 transporter | 37 | 11 | 41 | 58 | 47.44 | 84.06 | 4.76 | 4.75E-05 | Host-associated | Nonpigmented |
| **g003456**  peptide toxin processing or immunity | 32 | 9 | 46 | 60 | 41.03 | 86.96 | 4.64 | 0.0002 | Host-associated | Nonpigmented |
| **g004672_000001**  membrane transport system | 32 | 9 | 46 | 60 | 41.03 | 86.96 | 4.64 | 0.0002 | Host-associated | Nonpigmented |
| **g002149_1_000001** (mdaB)  NADPH: quinone oxidoreductase | 42 | 14 | 36 | 55 | 53.85 | 79.71 | 4.58 | 3.72E-05 | Host-associated | Nonpigmented |
| **g002851**  secreted serine protease involved in extracellular protein degradation | 36 | 11 | 42 | 58 | 46.15 | 84.06 | 4.52 | 9.07E-05 | Host-associated | Nonpigmented |
| **g001817** (speG)  spermidine N(1)-acetyltransferase | 36 | 11 | 42 | 58 | 46.15 | 84.06 | 4.52 | 9.07E-05 | Host-associated | Nonpigmented |
| **g002444_1_000001**  lipoprotein or antimicrobial compound transporter | 49 | 19 | 29 | 50 | 62.82 | 72.46 | 4.45 | 2.87E-05 | Host-associated | Nonpigmented |
| **g000237_1_000007** (endoI)  chitodextrinase | 65 | 37 | 13 | 32 | 83.33 | 46.38 | 4.32 | 0.0001 | Host-associated | Nonpigmented |
| **g000445**  L-2,4-diaminobutyric acid acetyltransferase | 51 | 21 | 27 | 48 | 65.38 | 69.57 | 4.32 | 3.14E-05 | Host-associated | Nonpigmented |
| **g001800**  periplasmic substrate-binding protein involved in quorum sensing | 51 | 21 | 27 | 48 | 65.38 | 69.57 | 4.32 | 3.14E-05 | Host-associated | Nonpigmented |
| **g005787_000002** (btuB)  vitamin B12 transporter | 39 | 13 | 39 | 56 | 50.00 | 81.16 | 4.31 | 0.0001 | Host-associated | Nonpigmented |
| **g002959_000001**  replication regulatory protein | 37 | 12 | 41 | 57 | 47.44 | 82.61 | 4.29 | 0.0001 | Host-associated | Nonpigmented |
| **g002567_000001** (nhaD)  Na(+)/H(+) antiporter | 53 | 23 | 25 | 46 | 67.95 | 66.67 | 4.24 | 3.35E-05 | Host-associated | Nonpigmented |
| **g002276**  structural component of type II or VI secretion system machinery | 49 | 20 | 29 | 49 | 62.82 | 71.01 | 4.14 | 6.15E-05 | Host-associated | Nonpigmented |
| **g002245**  membrane-bound O-acyltransferase | 40 | 14 | 38 | 55 | 51.28 | 79.71 | 4.14 | 0.0001 | Host-associated | Nonpigmented |
| **g003184_000002** (yybrR)  HTH-type transcriptional regulator | 43 | 16 | 35 | 53 | 55.13 | 76.81 | 4.07 | 9.72E-05 | Host-associated | Nonpigmented |
| **g000796_1_000001**  membrane-associated transcriptional regulator | 51 | 22 | 27 | 47 | 65.38 | 68.12 | 4.04 | 6.63E-05 | Host-associated | Nonpigmented |
| **g003378**  glutamate-dependent transcriptional regulation | 45 | 18 | 33 | 51 | 57.69 | 73.91 | 3.86 | 0.0001 | Host-associated | Nonpigmented |
| **g001818_1_000018** (baeR)  transcriptional regulatory protein | 46 | 19 | 32 | 50 | 58.97 | 72.46 | 3.78 | 0.0001 | Host-associated | Nonpigmented |
