## Supplementary Table 4 for "*Pseudoalteromonas* is a novel symbiont of marine invertebrates that exhibits broad patterns of phylosymbiosis"

**Supplementary Table 4: Metabolic functions and biogeochemical traits more commonly found in host-associated genomes.** A summary of the metabolic functions and enzymes associated with host-associated (t+) pigmented (orange) and nonpigmented (purple) genomes. The name of the enzyme or function is in bold, followed by a brief description of its function. The presence and absence of a function or gene is indicated by (g+) or (g-), respectively. An odds-ratio greater than one indicates that the gene/function is more commonly associated with the host-associated lifestyle strains.

| Function/Enzyme | g+t+ | g+t- | g-t+ | g-t- | sensitivity | specificity | Odds-Ratio | Fisher P | Lifestyle | Clade |
| --- | --- | --- | --- | --- | --- | --- | --- | --- | --- | --- |
| cellulase  cellulose degradation | 72 | 43 | 6 | 26 | 92.31 | 37.68 | 7.26 | 1.44E-05 | Host-associated | Pigmented |
| S09B  exopeptidase | 38 | 10 | 40 | 59 | 48.72 | 85.51 | 5.61 | 9.08E-06 | Host-associated | Pigmented |
| S01A  serine endopeptidase | 20 | 5 | 58 | 64 | 25.64 | 92.75 | 4.4 | 0.004 | Host-associated | Pigmented |
| Chitiniase  chitin degradation | 67 | 42 | 11 | 27 | 85.90 | 39.13 | 3.92 | 0.001 | Host-associated | Pigmented |
| GH018  chitinase  endo-N-β-D-glucosaminidase | 67 | 42 | 11 | 27 | 85.90 | 39.13 | 3.92 | 0.001 | Host-associated | Pigmented |
| GH020  glycoside hydrolase | 67 | 42 | 11 | 27 | 85.90 | 39.13 | 3.92 | 0.001 | Host-associated | Pigmented |
| Acetaldehyde => ethanol  ethanol fermentation | 30 | 11 | 48 | 58 | 38.46 | 84.06 | 3.30 | 0.003 | Host-associated | Pigmented |
| C56  endopeptidase | 63 | 15 | 3 | 7 | 95.45 | 31.82 | 9.80 | 0.002 | Host-associated | Nonpigmented |
