## Supplementary figures and images for "*Pseudoalteromonas* is a novel symbiont of marine invertebrates that exhibits broad patterns of phylosymbiosis"

### Extended Data Fig. 1

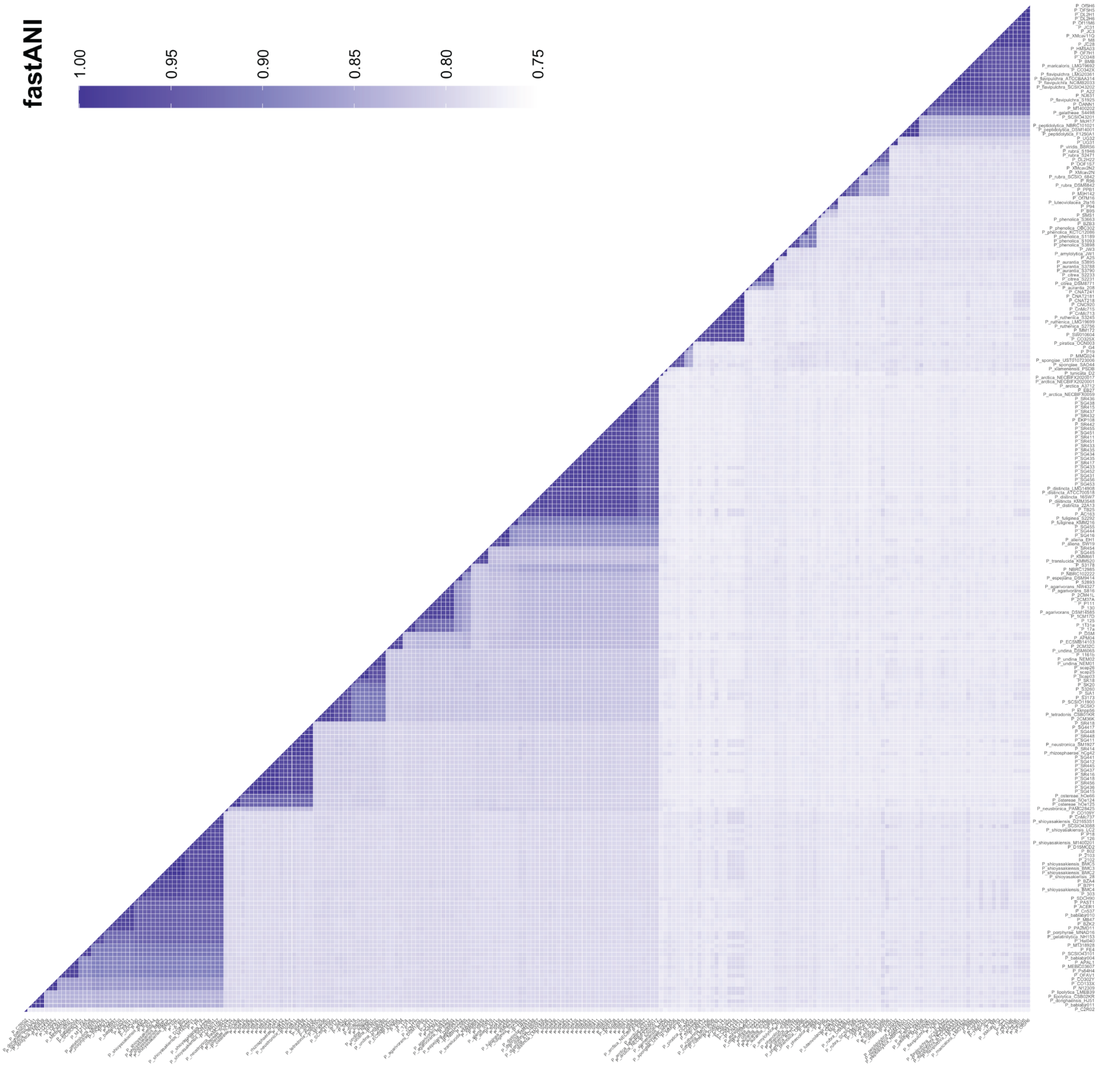

### Extended Data Fig. 2

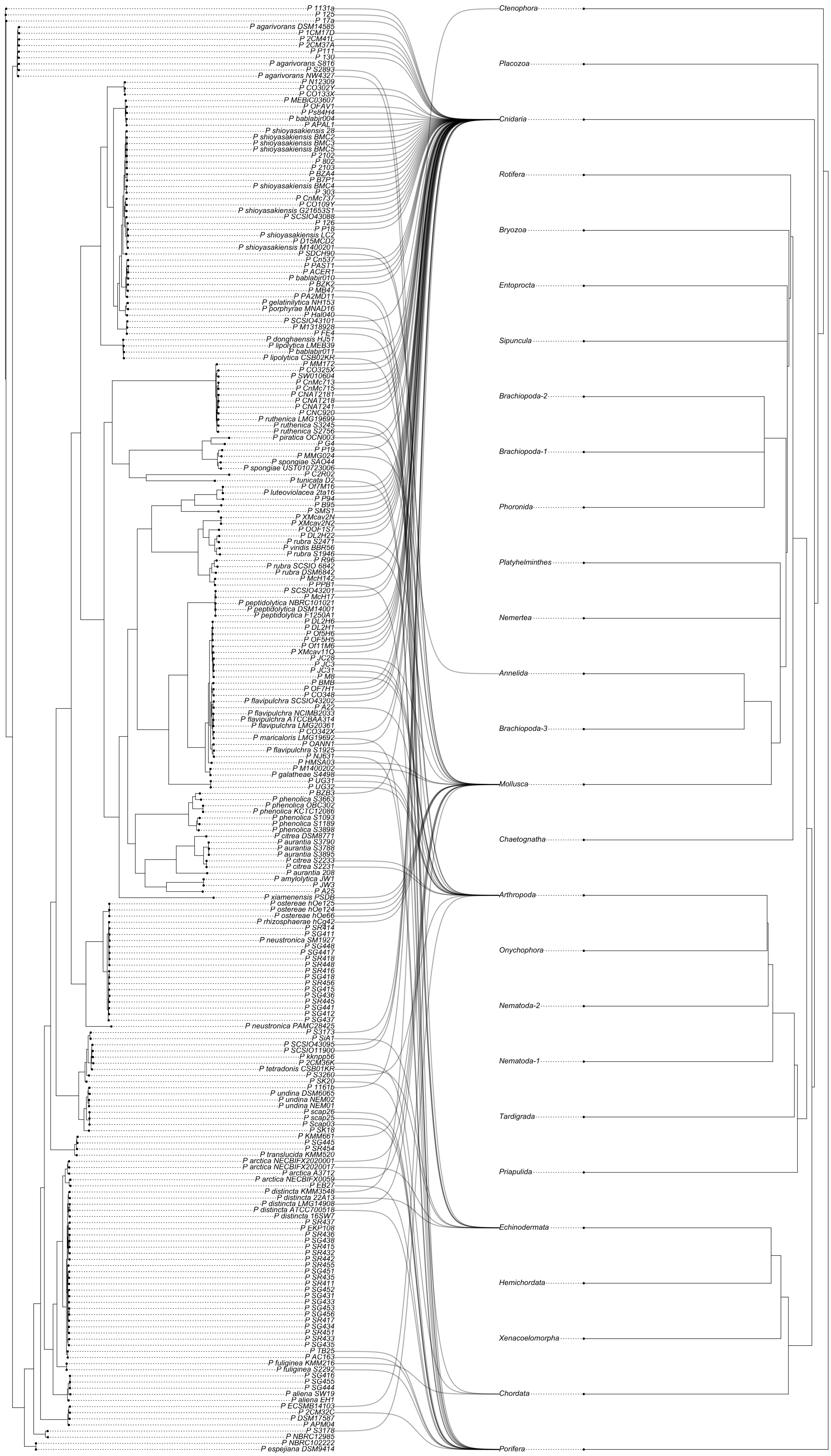

### Extended Data Fig. 3

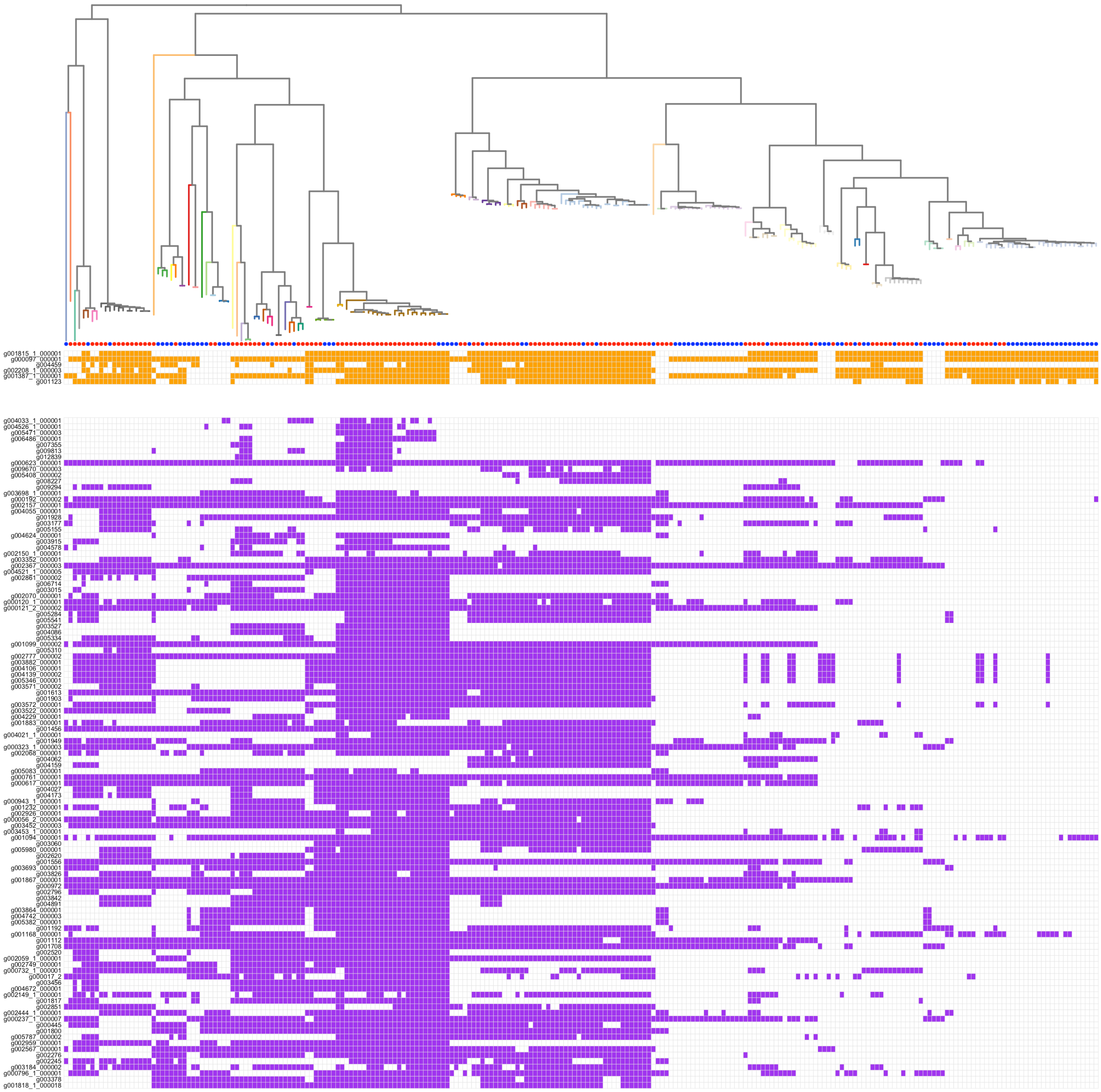
